# Real-time in vivo imaging of rodent brain response to hypoperfusion enabled by remotely controlled elastomeric micro-balloons

**DOI:** 10.1101/2025.09.03.673980

**Authors:** Jong Bin Kim, Jinghui Wang, Yinding Chi, Jingxian Wu, Alicia Ng, Guanda Qiao, Honglin Tan, Miroslaw Janowski, Piotr Walczak, Yajie Liang, Shu Yang

## Abstract

Current preclinical models of ischemic stroke in mice do not permit simultaneous and continuous in vivo brain imaging during the peri-stroke period, therefore missing critical pathophysiological events that could be pivotal for stroke management at the early stage. Here we report remote control of the blood flow of cerebral arteries in live mice continuously at different states to induce stroke in a precise, reliable, and reversible manner. The sub-millimeter scale micro-balloons can expand more than four times their initial diameter, featuring a monolithic elastomeric wall with selectively stiffened regions for controlled inflation and elasticity depending on the target vessels. By allowing for the control of common carotid artery diameter through a cuff pressing against the artery, the micro-balloon recapitulates clinically relevant haemodynamics in a mouse model of global brain ischemia, evidenced by real-time imaging of the brain through intravital microscopy and magnetic resonance imaging. The presented micro-balloons hold significant potential to improve the treatment of stroke patients for minimally invasive intervention and in vivo imaging of the pathophysiological events in the peri-stroke phase.

## Main

As one of the leading causes of death and long-term disability worldwide, ischemic stroke represents a major public health concern with substantial social and economic impacts^1^. It occurs when blood flow to the brain is reduced to the point where it can no longer meet the metabolic demands of neural tissue, leading to cellular injury and, if prolonged, irreversible damage, accounting for approximately 87% of all stroke cases^2,3^. In addition to the acute consequences, chronic cerebral hypoperfusion has been recognized as a significant risk factor for the development of cognitive decline and dementia^4,5^, further underscoring the critical importance of early diagnosis to understanding and modeling of impaired cerebral perfusion. Monitoring the brain’s immediate response to early ischemic stroke is crucial, as this hyperacute phase encompasses rapid pathophysiological events that significantly influence the extent of neuronal damage and the potential for recovery^6,7^. However, this critical window remains inaccessible for direct study due to significant delays^8–10^ between stroke onset and hospital admission of patients (**Fig. 1a**). Consequently, the early pathophysiological processes following stroke onset remain a “black box,” limiting our understanding and hindering the development of timely therapeutic interventions. Rodent models have been indispensable for uncovering mechanisms of ischemic stroke and testing therapeutic strategies^11,12^. Among them, mice are especially advantageous due to the availability of genetic tools that allow precise manipulation of specific molecular pathways involved in cerebrovascular injury and repair^13,14^. Nevertheless, most conventional approaches, such as middle cerebral artery occlusion or photothrombosis, lack the control of vascular occlusion in a remotely controlled manner under advanced in vivo imaging paradigms such as magnetic resonance imaging (MRI). Researchers have demonstrated feasibility of vascular recanalization and reperfusion through thrombolysis in a porcine model of ischemic stroke, using real-time MRI guidance^15^. However, real-time monitoring of the stroke onset remains impossible, limiting insights into the initial cerebrovascular and cellular responses. Moreover, the inability to precisely control the timing and repetition of occlusion events precludes systematic investigation of repeated or intermittent ischemia, which may better reflect clinical scenarios such as transient ischemic attacks or fluctuating perfusion in stroke-prone individuals^16,17^.

**Fig. 1.**
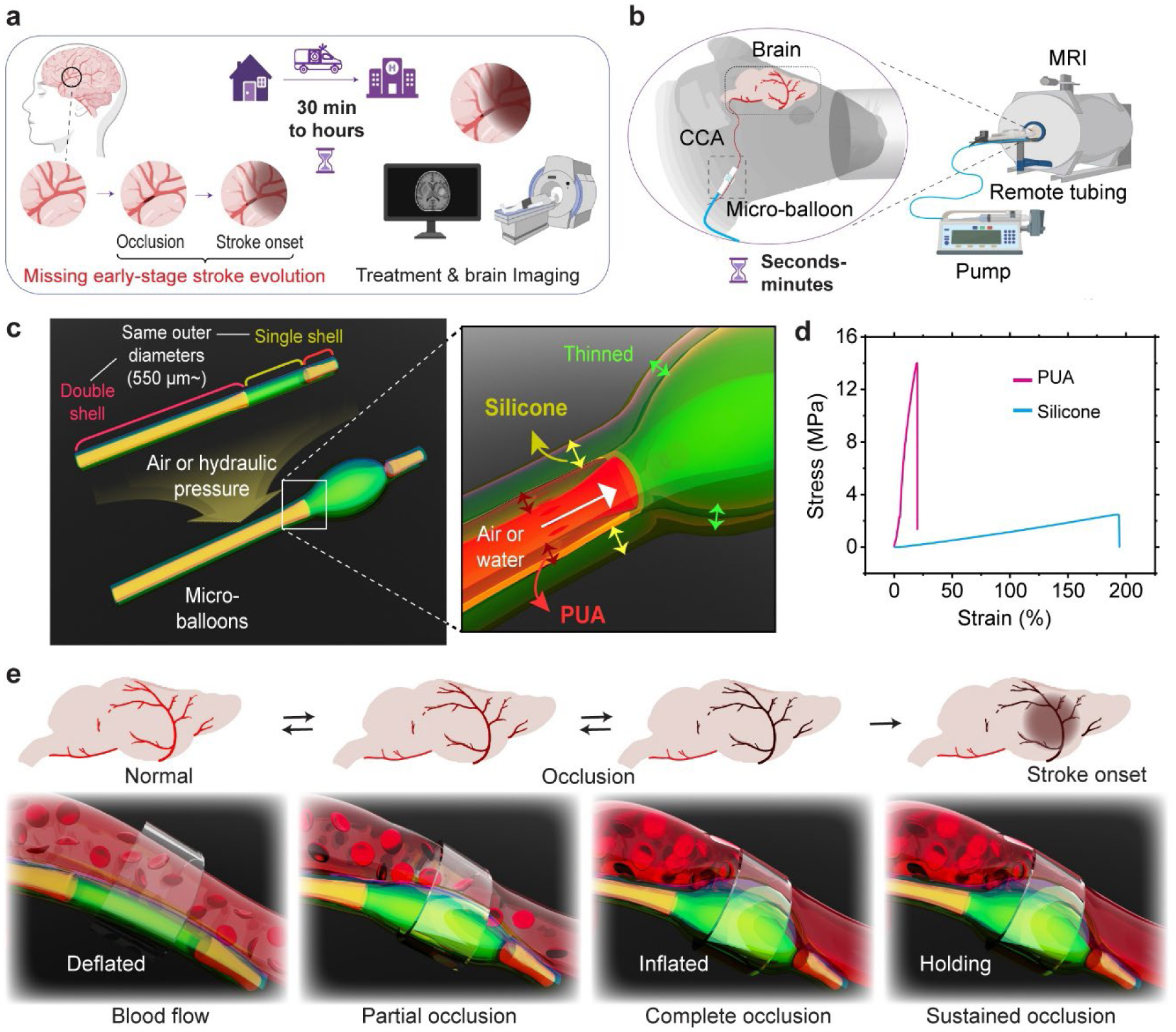
Concept and design of elastomeric micro-balloons for remote control of brain perfusion. **a,** Schematics illustrating the delay between stroke onset and hospital-based imaging and treatment for patients. **b,** Our strategy to image stroke onset in a mouse model using a remotely controlled, MRI-compatible micro-balloon. **c,** Schematics of the micro-balloon design, featuring inflation at a designated region with a magnified view. **d,** Stress-strain curves of the PUA and silicone. **e,** By controlling the inflation of the micro-balloon positioned adjacent to the common carotid artery within a confined space, the degree of occlusion can be precisely adjusted to model cerebral hypoperfusion or reperfusion.

Clearly, there is a significant gap in our understanding of how the brain responds to the onset of ischemia. Therefore, a micro-balloon that can reversibly change its diameter in a finely timed and repeatable manner offers a promising approach to understanding ischemic attacks by remotely blocking the blood vessel for the induction of ischemic events and the intermittent states. Soft robotic sleeves have been developed to modulate the size of the aorta for modeling aortic stenosis^18^, and millimeter-sized balloons are shown to induce aortic constrictions in small animal models^19^. However, these devices are neither small enough nor light enough to manipulate thin blood vessels such as those in mice. Clinically available balloon catheters can be compact with a diameter as small as 0.6 mm^20,21^, where a folded tubular balloon made from a non-stretchable polymer is joined with a metal-included shaft. However, its fabrication process is complicated, and folding limits further miniaturization of the balloon for different target vessels. Additionally, it cannot be expanded at intermediate states, nor return to the original state once unfolded. Most importantly, the inclusion of a metal component makes the catheter incompatible with MRI.

Here, we create highly tunable elastomeric micro-balloons that can be gradually inflated to block the blood vessel at different stages, remotely. The micro-balloons are MRI-compatible and small (550 μm to 900 μm in diameter) to target cerebral arteries to induce stroke in mice (**Fig. 1b**). The balloon is fabricated from a monolithic, straight silicone hollow tube, with certain regions of the inner wall coated with a thin layer of rigid polyurethane acrylate (PUA) (**Fig. 1c**). Upon pressurizing the entire tube, the non-coated region is selectively inflated into a balloon due to the large contrast in the elastic moduli: 1.3 MPa for silicone and 77.6 MPa for PUA (**Fig. 1d**). The inflated segment can maintain its shape at any intermediate state and recover rapidly and completely before it bursts. The micro-balloon is employed to dissect the effect of reversible occlusion of the common carotid artery on the blood flow, oxygenation level, and perfusion of the mouse brain through two-photon microscopy and MRI (**Fig. 1e**). The results will shed light on the brain’s response to the onset of ischemic stroke, which is hitherto an uncharted territory.

## Results

### Design and fabrication of the inflatable micro-balloons

Conventionally, centimeter-sized elastomeric balloons are fabricated by methods such as molding and sheet rolling^22,23^. Air injection into a tubular mold filled with uncured elastomer melt is relatively simple; however, the air core is typically off-centered due to the gravity effect^24^, leading to a tube with inconsistent thickness. Upon inflation, the tube bends, rotates, and twists, which is undesired for blood vessel occlusion. Here, we fabricated elastomeric micro-balloons of uniform thickness by air-jetting the precursors of desired rheological properties (**Fig. 2a**). First, microchannels (diameter, 550, 700, and 900 μm, respectively) were fabricated from elastomeric polydimethylsiloxane (PDMS), Sylgard 184, with Young’s modulus (*E*) of ∼1.8 MPa^25^, and used as the molds, which could be readily removed to release the micro-balloons. The PDMS precursor was poured over a long microneedle of different gauge sizes, which was removed after thermal curing. When the channel is rigid, the displacement of fluid by the injected air is described as the Bretherton’s problem^26,27^, where the flow of a long air bubble leaves a thin film along the wall due to the balance between the surface tension and the fluid resistance. Since our microchannels are elastic, and the displaced fluid is highly viscous (viscosity, ∼10^5^ Pa‧s) with a Capillary number (*Ca*) beyond the typical range for Bretherton’s problem, direct application of this model to our system is not feasible. Meanwhile, there is the airway reopening model^28,29^, where air propagation detaches and pushes fluid radially against an expanding channel without displacing it, preventing the formation of a hollow core after depressurization. In our system, because of its boxy geometry and relatively high elastic modulus, the expansion of the microchannels is limited. Therefore, a hollow core could be created after air injection, following the airway reopening model with the foundational principles observed in Bretherton’s problem.

**Fig. 2.**
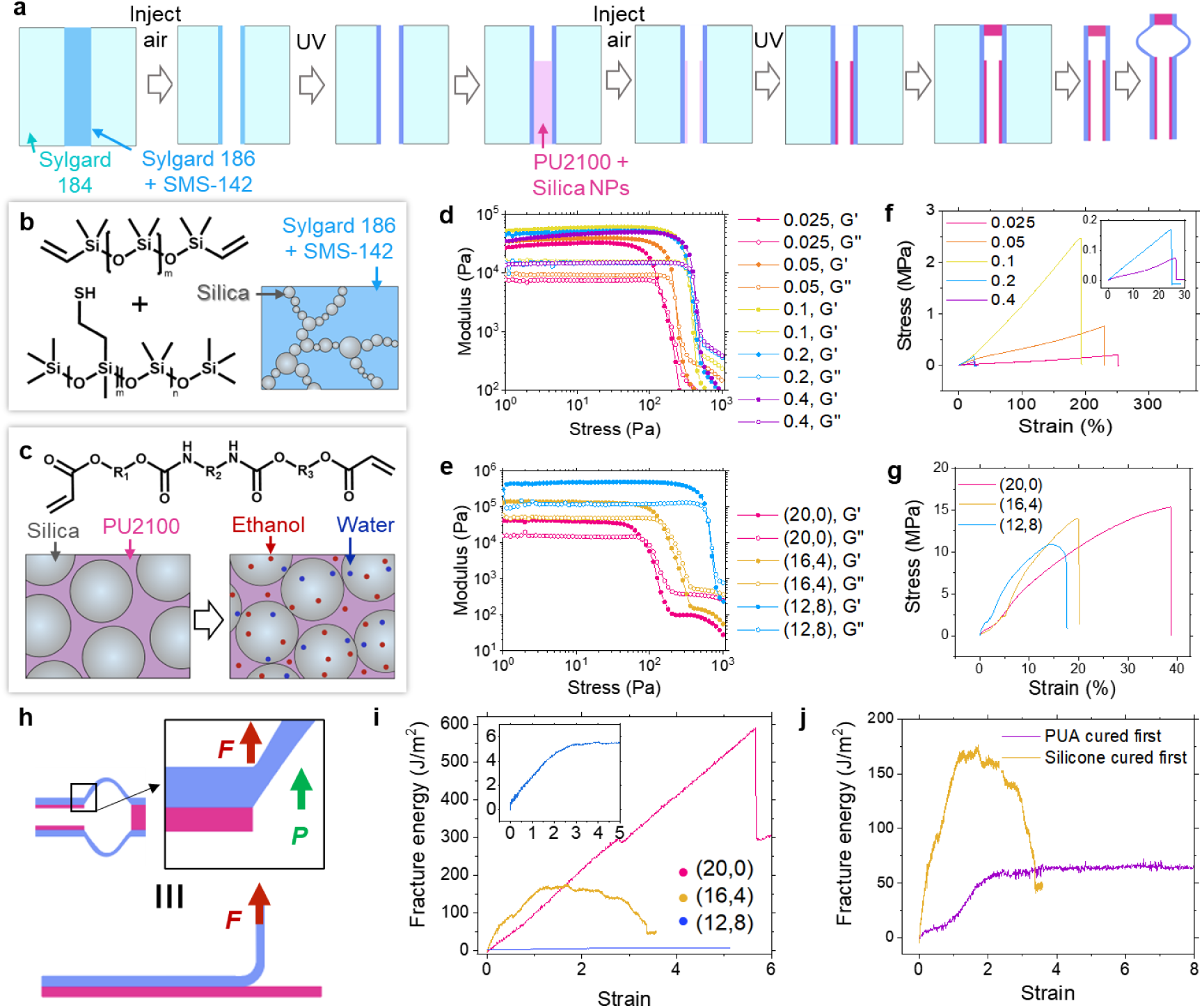
Fabrication process and properties of the consitutent materials. **a,** Schematics of the fabrication steps. **b,c,** Constituent components of the precursors for silicone (b) and PUA shells (c) with the illustrations of corresponding colloidal particle networks. **d–g,** Oscillatory tests (d,e) and tensile tests (f,g) of the precursors for silicone (d,f) and PUA (e,g) shells with different compositions. **h,** Schematic illustration of micro-balloon inflation, analogous to a peeling test. **i,** Adhesion forces between the silicone shell with m_=_/m_SH_ = 0.1 and PUA shells with different compositions. **j,** Adhesion forces with the curing order of the two shells, when the PUA shell has (16,4) solvent compositions.

The PDMS mold was then injected with a mixture (referred to as silicone precursor) consisting of a silicone precursor, Sylgard 186, hexamethyldisilazane (HMDS)-treated silica fillers, and PDMS copolymers with pendant mercapto groups, SMS-142, followed by air injection and UV curing to create a single-shell tube (see Experimental and Materials section). For a double-shell structure, a PUA precursor, PU2100 dispersed with silica nanoparticles (NPs), was injected into the single-shell tube, followed by air injection and UV curing (**Fig. 2a**). One end was left open for connection to a pneumatic system, and the other end was sealed to allow pressurization. After air injection, the air bubble and precursor are subject to instabilities inside the tube resulting from surface tension and gravity, leading to an undulated inner wall that would block the core. To address this, we employed yield-stress fluids (silicone and PUA precursors), which behave like solids below a critical stress and flow like liquids once exceeding the threshold.

The yield stress enables the shell structure to resist deformation caused by instability, maintaining its integrity unless a stress exceeding this threshold is applied. Specifically, a colloidal network is formed within the dispersion, establishing a percolated, space-spanning structure that resists deformation at a low stress (**Extended Data Fig. 1**). By tailoring each precursor’s mechanical properties after curing, Silicone and PUA behaved linearly elastic at low shear rates (**Extended Data Fig. 2**)^30,31^. When the silica nanofillers in the precursor percolate, the apparent yield stress reaches 400 Pa (**Fig. 2b** and **Extended Data Fig. 2a**)^32,33^. We note when silica NPs (diameter, 250 nm) are first introduced into the PUA precursor, the presence of the acrylate layer around the NP surface makes them repel each other (**Fig. 2c**)^34^. Upon adding trace solvents—ethanol and water, which disrupt the acrylate layers due to the solvents’ high affinity for silica, silica NPs become attractive and form a network, yielding a 3.5-fold increase in *E* (77.6 MPa) over that of the crosslinked pure PUA (19.4 MPa; **Extended Data Fig. 3a**).

We next examined how precursor formulations affect rheological properties. Yield stress must be high enough to resist the instabilities, yet low enough to allow it to be displaced by air injection. We varied the molar ratios of Sylgard 186 to SMS-142, that is vinyl (m_=_) to thiol groups (m_SH_), from 0.025 to 0.4. As seen in **Fig. 2d**, the highest storage modulus, *G*”, was achieved when m_=_/m_SH_ = 0.1, indicating an enhanced elastic response and reduced susceptibility to instabilities. Given that it also exhibited a relatively high apparent yield stress, this formulation was suitable for maintaining structural stability (**Extended Data Fig. 2a**). Although a higher viscosity also increased flow resistance, the increase was on the same order as that by varying m_=_/m_SH_, suggesting that it is not a dominant factor for producing well-defined shells (**Extended Data Fig. 2c**). For the PUA precursor, addition of water increased yield stress more significantly than that of ethanol due to its preferential interactions with silica over acrylate. As seen in **Fig. 2e**, **Extended Data Fig. 2b**, and **2d**, the weight fractions of ethanol (16 wt%) and water (4 wt%) relative to that of silica NPs, referred to as (16,4), were optimal. The rheological properties of the chosen compositions of silicone and PUA precursors are shown in **Extended Data Fig. 4**: *G*” = ∼135 kPa and 60 kPa for the PUA and silicone shells, respectively, and the apparent yield stresses and viscosities for both shells are ∼ 400 Pa and ∼10^5^ Pa‧s at 0.0001 s^-1^, respectively. The *G*” and apparent yield stresses are sufficiently high to confer a solid-like property, yet the viscosities remain low enough to minimize flow resistance. This balance of rheological properties facilitates effective air propagation and precursor displacement in our tubes, which will be discussed later.

### Mechanical properties for efficient micro-balloon functionality

The mechanical properties of both crosslinked shells are crucial for the performance of the micro-balloons. Tensile testing of the silicone shell shows that at m_=_/m_SH_ = 0.1, silicone with *E* ∼ 1.3 MPa can extend by three times, which prevents the tube from tearing during demolding (**Fig. 2f**). When SMS-142 is in excess (m_=_/m_SH_ = 0.1), the SH-rich side chains of SMS-142 promote dense and efficient network formation, while Sylgard 186 becomes the limiting reactant. Because silicone is isotropic, its shell exhibits mild hysteresis under cyclic loading at 60% strain, which is sufficient for blood vessel occlusion (**Extended Data Fig. 3b**). PUAs crosslinked from precursors with ethanol/water weight ratios of (20, 0), (16, 4), and (12, 8) had *E* = 58.8, 77.6, and 113.0 MPa, respectively, which are much higher than that of silicone. Therefore, the optimal formulation of a PUA shell should be guided by rheological behaviors rather than tensile properties (**Fig. 2g**).

Adhesion between the silicone/PUA shells during inflation is critical. The unreacted thiols in the crosslinked silicone form covalent bonds with the newly cast PUA under UV^35^, which is confirmed by a peeling test (**Fig. 2h**). The maximum fracture energies for PUA prepared from (20, 0), (16, 4), and (12, 8) ethanol/water compositions are 591.9, 169.0, and 5.4 J/m², respectively. This trend can be attributed to the hydrophobicity of the silicone shell, as water in PUA precursors increases interfacial energy with silicone during fabrication. Thus, increased water content in PUA markedly reduces adhesion (**Fig. 2i)**. To confirm this, we blade-coated the silicone precursor and briefly cured it for 5 s, followed by application of the PUA precursor and exposure to UV light to form a covalent bond between them. Although the (20, 0) composition shows the strongest adhesion among the three formulations, it lacks the yield stress required for fabrication of the shells (**Fig. 2e**). In comparison, the (16, 4) composition has a fracture energy sufficiently high to prevent delamination, as shown later. The hypothesis of covalent-bond-assisted adhesion was supported by a control experiment, where the PUA shell was cured first, yielding ∼3× lower adhesion than that when silicone was cured first (**Fig. 2j**).

### Optimization of the number of balloon shells and the thickness

To achieve the desired expansion and force generation, the elastomer shell thickness must be carefully tailored. The silicone precursor was deposited once or twice onto the inner wall of the PDMS microchannels (diameters, 550, 700, and 900 μm), followed by air injection, and selective PUA deposition (**Fig. 3a**). The longitudinal uniformity was achieved when air was injected at a constant air front speed, in line with Bretherton’s equation (**Fig. 3b** and **Extended Data Fig. 5, 6**)^27^. In contrast, the shell thickness increased along the channel when air was injected under a constant pressure because the fluid-filled, flow-resisting region was shortened as the air front speed gradually increased (**Extended Data Fig. 7**). While Bretherton’s model— and its extensions, such as the yield stress fluid correction by Laborie *et al.*^36^—can calculate the shell thickness in rigid channels, predicting thickness in elastomeric microchannels is challenging since the wall inflates during air injection. Nonetheless, the same speed-dependent trend empirically defines a processing window for a target thicknesses (**Fig. 3c**). When a second elastomer shell was applied, thickness variation across the channel sizes was significantly reduced (**Fig. 3d**). This is likely because the initial channel size difference is compensated in the first shell deposition, reducing structural dependence on channel size in the subsequent shell deposition.

**Fig. 3.**
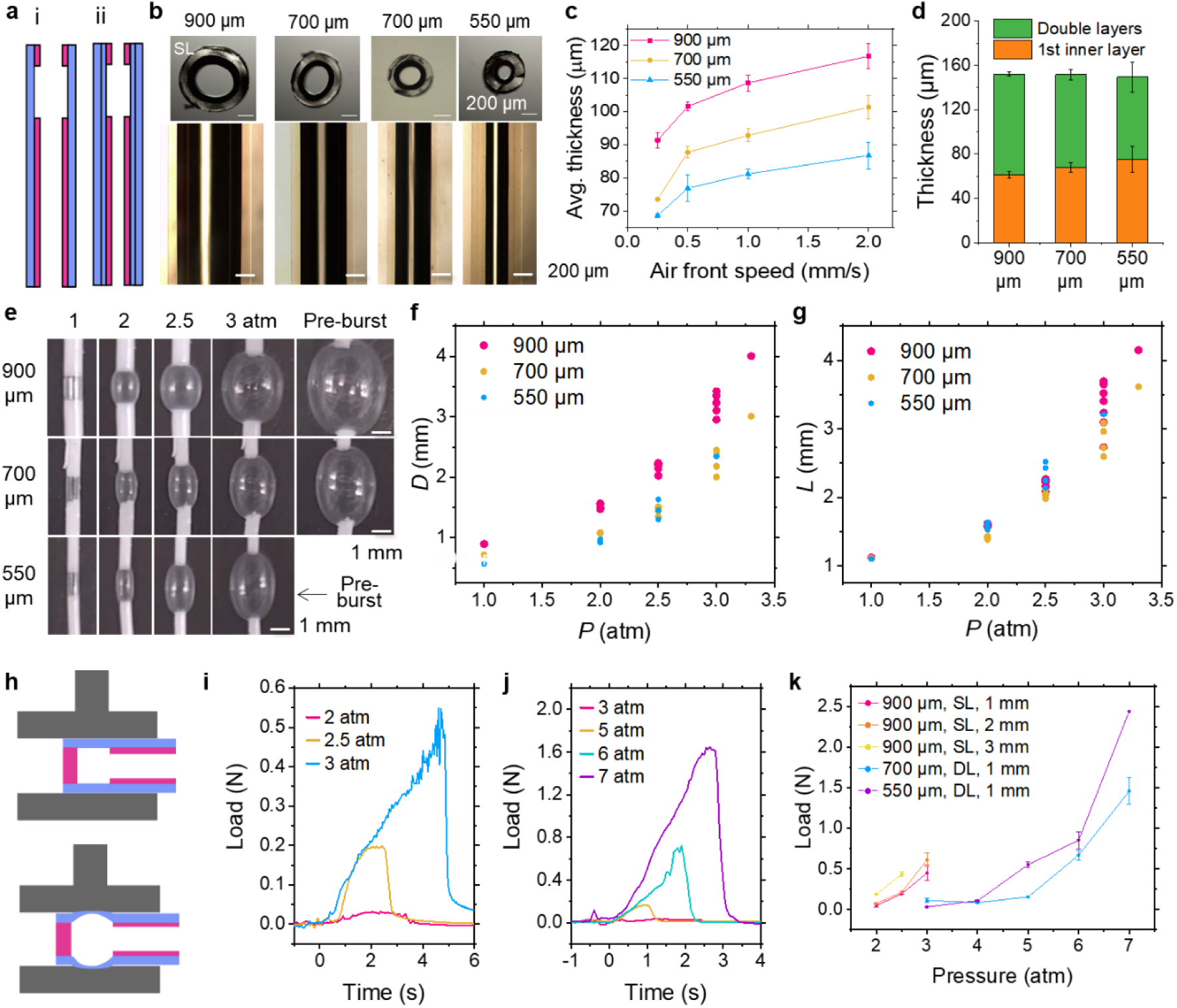
Pneumatic inflation behaviors of micro-balloons with different designs. **a,** Schematics and the cross-sectional views of micro-balloons. **b,** Cross-sectional optical microscopy (OM) images of the micro-balloons comprising both shells (top) and their top views composed of the silicone shell only (bottom) at various outer diameters. **c,** Average thicknesses of the elastomer shell as a function of air front speed (*n* = 3). **d,** Change in the average thickness after two sequential cycles of silicone precursor filling and air injection at 0.5 mm/s (*n* = 3). **e,** Digital photos of the micro-balloons at different pressures. **f,g,** Pressure-dependent changes in the diameter (f) and length (g) of balloon segments (*n* = 3). **h,** Schematics of the expansion force measurement setup by a universal testing machine. **i–k,** Expansion forces of the 20G micro-balloons as a function of pressure for a single shell (i), double shell (j), and varying dimensions and shell counts (k) (*n* = 3).

### Pneumatic Response of Micro-Balloon Tubes

The micro-balloon with a single elastomer layer was gradually inflated to four times its original size with increasing pneumatic pressure at ∼3 atm (**Fig. 3e**), whereas those with thicker, double shells expanded less than twice their original size (**Extended Data Fig. 8**). Both single- and double-shell micro-balloons undergo not only inflation but elongation during pressurization. For a 1 mm-long balloon, the inflated length (*L*) and maximum diameter (*D*) across various initial diameters fall within the pressure-dependent operation window (**Fig. 3f,g**). The aspect ratio of *L*/*D* decreases with the increase of pressure, indicating that lateral expansion becomes more pronounced at higher inflation levels (**Extended Data Fig. 9**). As the strain increased, irreversible deformation occurred, reflecting the viscoelastic nature of the shell materials. As a result, the balloons might stretch more at the same pressure during subsequent inflations, particularly at 3 atm for single-shell balloons. Importantly, the inflation and contraction are reversible within ∼2 atm, at which the balloon expands approximately twice its diameter, sufficient to occlude a blood vessel. This unique capability sets our balloon apart from the clinical catheters. In addition, injection of water instead of air could allow more precise control of balloon inflation at precise stages. Lastly, as balloon length increased, the pressure required to enlarge the lateral diameter decreased, while the maximum achievable diameter remained more or less the same (**Extended Data Fig. 10**).

Since the force exerted during expansion emerges as another key characteristic of the balloons, we measured the expansion forces using a universal testing machine (**Fig. 3h** and **Extended Data Fig. 11**). A single-shell micro-balloon with a 900 μm outer diameter exerted 0.03, 0.20, and 0.55 N at 2, 2.5, and 3 atm, respectively (**Fig. 3i**). Double-shell micro-balloons sustained higher pressure and exerted greater forces, 0.17, 0.71, and 1.65 N at 5, 6, and 7 atm, respectively (**Fig. 3j**). The expansion force increased with the initial balloon length. At a given pressure, smaller-diameter tubes exerted higher forces since thinner walls are less resistant to expansion (**Fig. 3k**). For instance, a double-shell micro-balloon (700 μm in diameter, 1 mm-long) exerted a maximum force of 2.5 N, making it suitable for applications such as the creation of occlusion for modeling stroke, expansion of calcified or highly resistant plaques in blood vessels, or opening resistant vasospasm.

### Remote-Controlled and Tunable Micro-Balloon System Enables Reliable In Vivo Vascular Occlusion

To evaluate the reliability and tunability of our device for remote vascular compression, we conducted in vivo tests to occlude the common carotid artery (CCA) of a mouse (**Fig. 4a**), by real-time monitoring of the cerebral blood flow rate in the brain through laser Doppler flowmetry. We first assembled the remote injector system filled with water for hydraulic control of the degree of expansion of the micro-balloon (**Extended Data Fig. 12a**). To provide a spatially constrained space, a customized C-shape plastic cuff was wrapped around both the micro-balloon and the target vessel (**Extended Data Fig. 12b**). The expansion of the micro-balloon was found positively correlated with the volume of water injected into the tubing (**Extended Data Fig. 13**). Gradual inflation and deflation of the micro-balloon through remotely controlling the position of a syringe plunger is highly desired for blocking or unblocking the CCA on demand (see **Fig. 4b**). Importantly, our remote-controlled platform enables stable holding at any of the expansion states lasting up to 10–20 s (but not limited to). No change in cerebral blood flow rate was observed when 1/3 stop-volume (1/3 v) was dispensed by the syringe. A significant decrease of flow from 536.9 perfusion unit (PU) to 410.1 PU (23.6%) occurred upon dispensing 2/3 v (**Fig. 4c**). Cerebral blood flow rate dropped by 96.5% (from the mean flux of 546.5 to a mean flux of 18.8, **Fig. 4c**) when the CCA was completely occluded (1 v, **Fig. 4c**). This was followed by a hyperperfusion response evidenced by the significantly higher flow rate when the injection volume is reduced from 1 v to 2/3 v (**Fig. 4c**). These changes were supported by statistics from 3 mice (**Fig. 4d**): open vs. partial open, *p* = 0.001, *n* = 11 trials; open vs. fully occluded, *p* = 0.0005, *n* = 12; open vs. reopened, *p* = 0.0093, *n* = 12. Interestingly, hyperperfusion was not observed when reopening was performed in a non-step manner (*p* = 0.0625, *n* = 5, **Fig. 4e**).

**Fig. 4.**
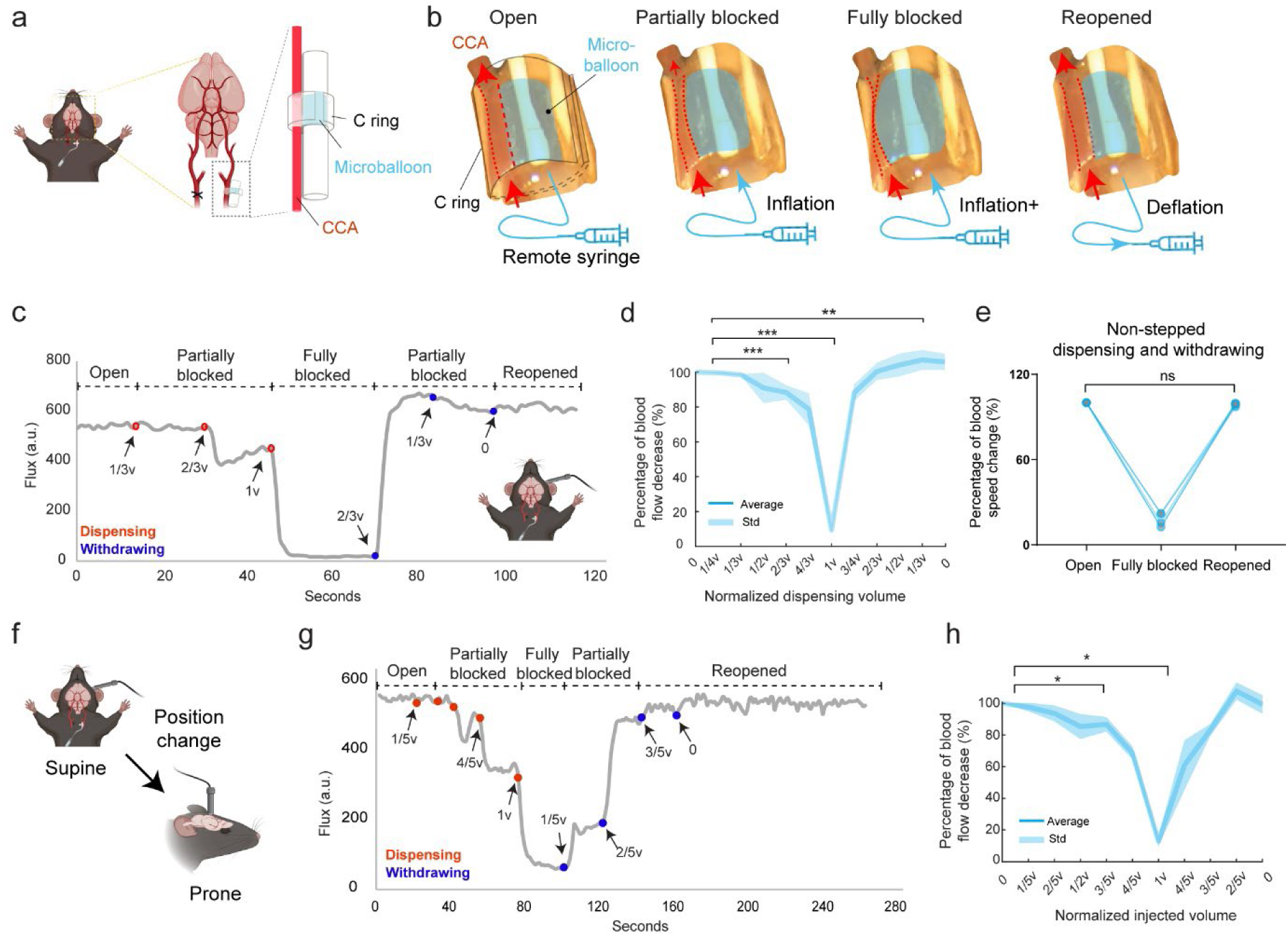
Laser Doppler recorded blood flow rate in the mouse brain upon manipulation of CCA occlusion at different levels. **a,** Surgical setting and micro-balloon placement on one side of the CCA. **b,** Vascular shape changes under non-inflated, inflated, and deflated conditions (indicated by red dashes). **c, d,** Blood flow rate recorded in the supine position (*n* = 12 trails from 3 mice; opened vs. partially blocked, *n* = 11 trials from 3 mice). **e,** Quantification of non-step manners (*n* = 5 trails from 3 mice). **f–h,** Blood flow rate recorded in the prone position (*n* = 6 trials from 4 mice). For normalization, the volume of liquid injected into the micro-balloon was defined as a single stop-volume (1 v). Differences were analyzed by Wilcoxon matched pairs signed rank test, ns, not significant, *\*p <* 0.05, *\*\*p <* 0.01, *\*\*\*p <* 0.001.

Since most in vivo imaging studies are conducted with mice in the prone position, we assessed the robustness of micro-balloon occlusion of the CCA after gently flipping the mice from supine to prone position. As the CCA was gradually occluded, a clear trend of decreasing blood flow velocity was observed (**Fig. 4f**). Partial occlusion to 4/5 v resulted in a 37.07% reduction in flux (from 550.9 to 346.72, **Fig. 4g**), and further occlusion to 1 v causes an 83.7% reduction in flow velocity (flux from 572.9 to 93.5). Statistics revealed significant difference between the open and fully blocked phases from 4 mice (*p* = 0.0312, *n* = 6), or the open and partial occlusion phases (*p* = 0.0312, *n* = 6, **Fig. 4h**). However, no significant decrease was observed between the open and reopened conditions (*p* = 0.8438, *n* = 6). Clearly, our remote micro-balloon actuating system is robust and yet tunable for in vivo blood vessel occlusion at different stages under either supine or prone position.

### Dynamic Cerebral Hemodynamics and Vessel Reactivity Monitored by High-Resolution Intravital Two-Photon Microscopy

Next, we leveraged an intravital two-photon fluorescence microscope (TPFM) to quantify real-time changes in cerebral blood flow rate during CCA occlusion. Line-scan under two-photon excitation permits high-speed tracking of the blood flow rate up to 84 mm/s in the brain^37,38^ and was performed over large blood vessels on the dorsal surface of the living mouse brain at 1 kHz (**Fig. 5a**). Inflation of the micro-balloon to the stop volume resulted in a marked reduction in flow rate (89.01%, from 6.05 to 0.66 mm/s, **Fig. 5b**), supported by statistics from 3 mice (open vs. blocked, *p* = 0.0005, *n* = 12 blood vessels, **Fig. 5c**). Flow rate rapidly recovered upon balloon deflation (blocked vs. re-opened, *p* = 0.0005, *n* = 12). There was no significant difference between the open and re-opened ones in terms of the blood flow rate (*p* = 0.5693, *n* = 12). More examples of line-scan plots shown in **Extended Data Fig. 14** reveal the acute and reversible hemodynamic impact of the targeted vascular occlusion with high temporal precision.

**Fig. 5.**
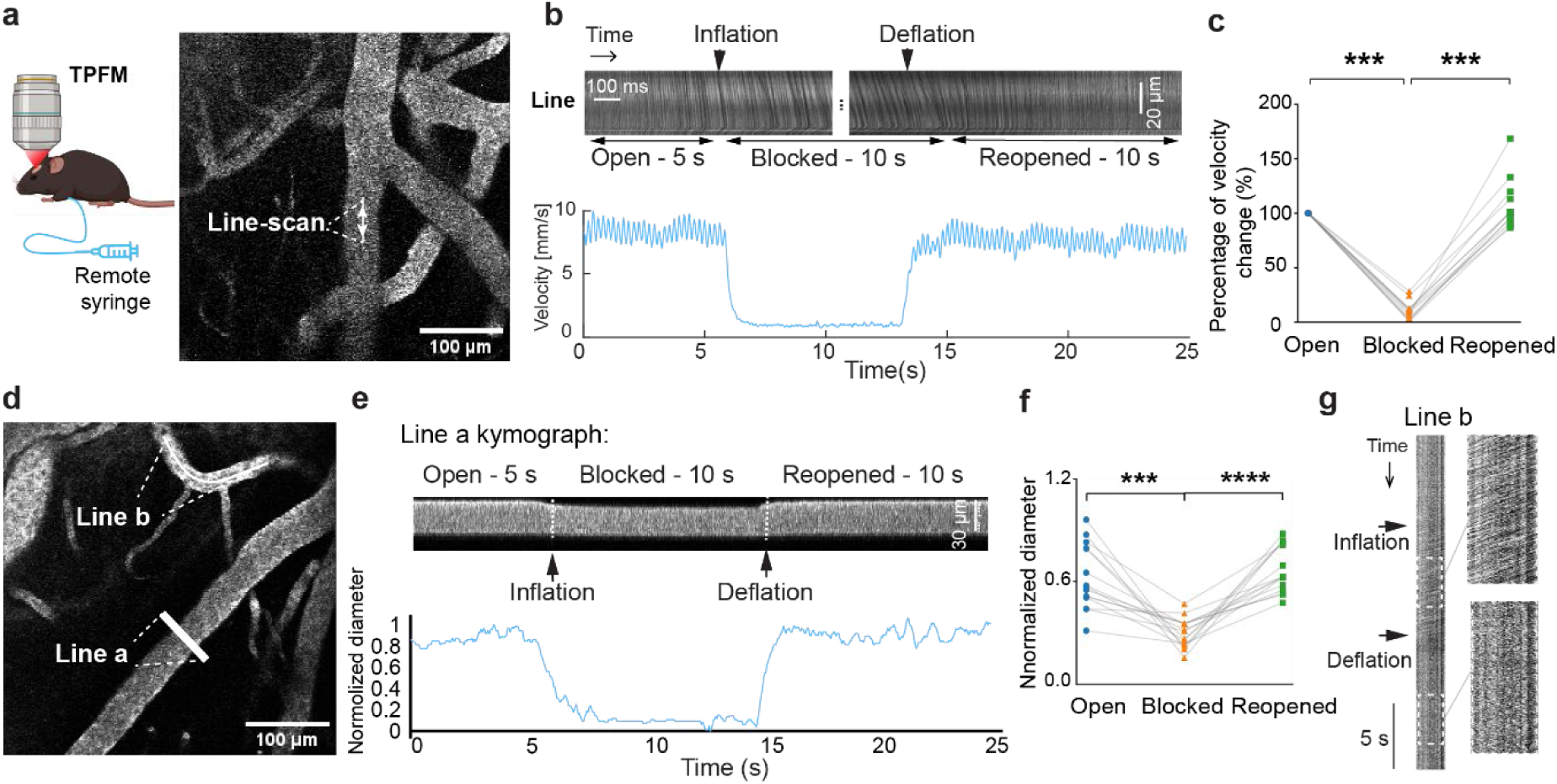
Intravital two-photon imaging of the mouse brain perfusion in response to micro-ballon manipulation of CCA. **a,** Example image of cortical vasculature of a mouse with a cranial window using TPFM. The line marks the locations of line-scan data acquisition. **b,** Space-time image from raw line-scan data (upper panel) and line-scanning particle image velocimetry (LS-PIV) analysis of flow rate over time (lower panel). **c,** Quantification of the flow rate of blood vessels (*n*=12 lines from 3 mice). Differences were analyzed by the Wilcoxon matched pairs signed rank test, *\*\*\*p <* 0.001. **d,** Example image of cortical vasculature using Two-photon imaging blood flow at 58 Hz frame rate. **e,** Kymograph for Line a show the change of flow upon micro-balloon manipulation. **f,** Quantification of blood vessel diameter changes (*n* = 14 vessels from 3 mice). Statistical analysis was performed using a paired t-test. *\*\*\*p =* 0.0001, *\*\*\*\*p<* 0.0001. **g,** Kymograph for Line b shows flow rate change upon blockage of CCA evidenced by the appearance of angled streaks.

Because line-scan imaging is limited to one-dimensional (1D) velocity measurements and cannot capture concurrent morphological changes (e.g., vessel dilation or constriction) during occlusion, we resorted to full-field raster scanning of cortical blood vessels at a fast frame rate (58 Hz) (**Fig. 5d)**. Interestingly, a rapid vasoconstrictive response upon complete occlusion was observed (**Fig. 5e**), which was reversed immediately following reperfusion. This transient constriction was supported by statistics from 3 mice (*p* = 0.0001, *n* = 14 blood vessels, **Fig. 5f**). More examples of kymographs are shown in **Extended Data Fig. 15**. The kymograph of the segmented line over a blood vessel in the two-dimensional (2D) time-lapse stack also revealed the drastic reduction in flow rate evidenced by the angled streaks after inflation of the balloon (**Fig. 5g**). This suggests 2D raster scanning under TPFM could provide both speed and morphology changes of the blood vessel in response to CCA blockage. Collectively, these results validate the micro-balloon’s efficacy for precise, reversible induction of cerebral ischemia in vivo. Using high-speed two-photon imaging, we combined 1D line-scans for quantitative flow with 2D raster scans for morphology. This approach revealed rapid vascular responses—flow arrest and transient vasoconstriction—that illuminate the mechanisms of hypoperfusion and reperfusion in stroke.

### Real-Time Mapping of Cerebral Hypoperfusion and Ischemic Damage Using Multi-Modal MRI in a Micro-Balloon Occlusion Model

To evaluate hemodynamic and oxygenation changes across diverse brain regions during micro-balloon-induced CCA occlusion in mice, we applied multi-modal MRI (**Fig. 6a**). Blood oxygen level-dependent (BOLD) MRI captured real-time tissue oxygenation shifts via deoxyhemoglobin variations under hypoperfusion. Balloon inflation triggered rapid BOLD signal reductions in both hemispheres (**Fig. 6b–d**), with swift recovery upon deflation. Regional variations were evident, with pronounced drops in contralateral cortex from 3 mice (same to the studies reported below), 18.4 ± 4.43% reduction, 95% CI: 13.7–23%, *p* = 0.0002, *n* = 6 tests (blue traces in **Fig. 6d**), followed by hippocampus, 9.96 ± 4.44% reduction, 95% CI: 5.3–14.6%, *p* = 0.0027, *n* = 6 (red traces in **Fig. 6d**) and dorsal striatum, 11.6 ± 4.24% reduction, 95% CI: 7.81–16.1%, *p* = 0.0312, *n* = 6 (green traces in **Fig. 6d**), and minimal in vertebrobasilar regions such as ventral striatum, 0.487 ± 0.346% reduction, 95% CI: −0.372–1.35%, *p* = 0.25, *n* = 3 (black traces in **Fig. 6d**). Ipsilateral sides also showed significant changes for these brain regions (cortex: 16.1 ± 4.41% reduction, 95% CI: 11.5–20.8.6%, *p* = 0.0003, *n* = 6; hippocampus: 8.34 ± 2.43% reduction, 95% CI: 5.79–10.9%, *p* = 0.0003, *n* = 6; striatum: 9.18 ± 2% reduction, 95% CI: 7.08–11.3%, *p* = 0.0312, *n* = 6; **Fig. 6d,e**). The change is more obvious in contralateral side compared to ipsilateral side (cortex: *p* = 0.0047, *n* = 6; hippocampus: *p* = 0.1156, *n* = 6; dorsal striatum: *p* = 0.0312, *n* = 6, **Fig. 6e**). Standard deviation (STD) maps (**Fig. 6b,c**, bottom row) highlighted the regional heterogeneity, which can be attributed to varying contributions from the vertebrobasilar arterial system. BOLD MRI dynamics from more animals could be found in **Extended data 16**.

**Fig. 6.**
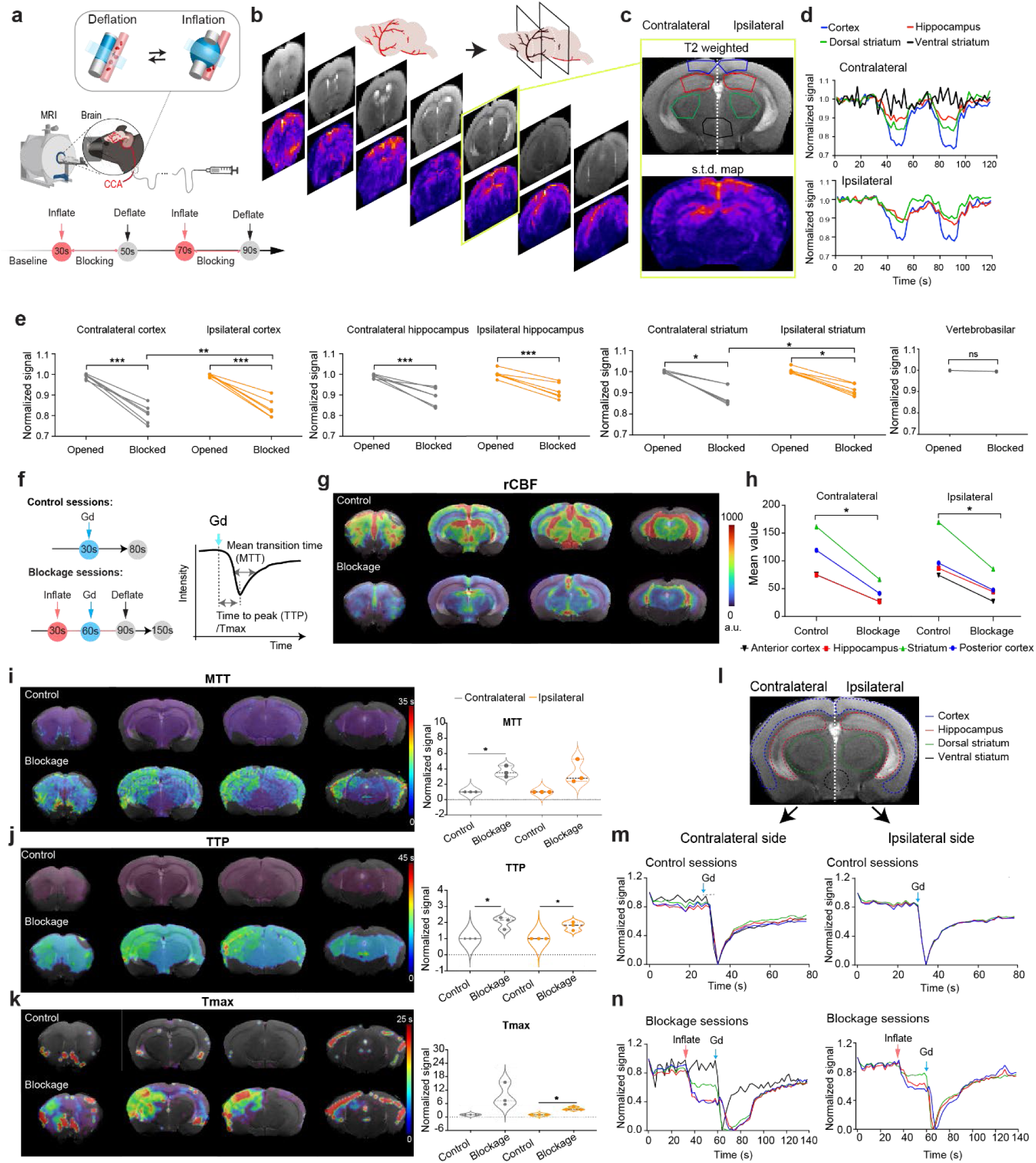
MRI study before, during, and after CCA blockage via micro-balloon. **a,** Schematic and experimental timeline for micro-balloon occlusion of the CCA. **b,** Representative coronal T2-weighted images (top) and corresponding oxygenation s.t.d. maps (bottom) during CCA occlusion. **c,** The example coronal plane for quantification and corresponding s.t.d. map. **d,** Normalized oxygenation signal time courses from different brain regions from ipsilateral and contralateral brain. **e,** Quantification of normalized oxygenation signals from 3 mice, in cortex (*n* = 6), hippocampus (*n* = 6), striatum (*n* = 6), and vertebrobasilar system (*n* = 3). *\*p <* 0.05, *\*\*\*p <* 0.001, paired t-test. **f,** Timeline of Gd-enhanced T2-weighted MRI before and during CCA blockage using a micro-balloon. **g,** Representative rCBF maps reconstructed from T2-weighted MRI before (control sessions) and during CCA blockage (blockage session). Gd-DTPA was administered via tail vein injection at indicated time points. **h,** Quantification of rCBF values in the posterior cortex, hippocampus, striatum, and anterior cortex, comparing contralateral and ipsilateral hemispheres. *\*p <* 0.05, paired t-test. **i–k,** Representative maps and quantification of MTT, TTP, and Tmax from T2-weighted MRI before and after micro-balloon-induced CCA blockage. Violin plots compare values from contralateral and ipsilateral hemispheres at baseline and during blockage. *n* =3, *\* p <* 0.05, one-sample t-test. **l–n,** Traces of signals in different brain regions (cortex, hippocampus, striatum, and vertebrobasilar system) are shown before (m) and during (n) CCA blockage.

Complementing BOLD, dynamic susceptibility contrast (DSC) MRI with gadolinium (Gd) was performed to quantify perfusion dynamics in control or CCA blockage conditions (**Fig. 6f**). Relative cerebral blood flow (rCBF) maps revealed marked ipsilateral reductions after occlusion (**Fig. 6g**) across regions (**Fig. 6h**; contralateral: *p* = 0.01, *n* = 3 regions; ipsilateral *p* = 0.0104, *n* = 3). Parametric maps showed baseline symmetry in mean transit time (MTT), time to peak (TTP), and time to maximum (Tmax) (top left panels in **Fig. 6i–k**, respectively), respectively, shifting to significant elevations after blockage (bottom left panels in **Fig. 6i–k**, respectively). Normalized signals increased ∼5-fold (MTT, **Fig. 6i**, right panel), ∼3-fold (TTP, **Fig. 6j**, right panel), and ∼6-fold (Tmax, **Fig. 6k**, right panel) contralaterally during occlusion (MTT: *p* = 0.0245; TTP: *p* = 0.0456; Tmax: *p* = 0.1081 *n* = 3 mice). Ipsilateral side exhibited similar significant delays in perfusion dynamics (MTT: *p* = 0.1097; TTP: *p* = 0.0348; Tmax: *p* = 0.0192; *n* = 3). Example DSC-MRI perfusion dynamics from more animals could be found in **Extended data 17.** Detailed analysis of DSC signal-time curves reinforced these findings (**Fig. 6l–n**). Baseline curves without blockage from different brain regions unanimously showed steep Gd-induced drops and quick recovery (**Fig. 6m**). Inflation of the micro-balloon induced delayed peak arrival (∼10–20 s shift), shallower minima, and prolonged transit, most severe in cortex/hippocampus, moderate in dorsal striatum, negligible in ventral striatum (**Fig. 6n**, left panel). These changes are more pronounced in the contralateral side of the brain (**Fig. 6n**). These findings align with parametric elevations and underscore reversible ischemic gradients via collateral flow.

## Discussion

We developed micro-balloons for remotely controlled, reversible vascular occlusion in mouse models of ischemic stroke, enabling compatibility with advanced biomedical imaging modalities such as MRI and two-photon microscopy. Specifically, we i) miniaturized the balloons to sub-millimeter scales (550–900 μm diameter) for targeting cerebral arteries of mice, ii) ensured MRI compatibility by eliminating metallic components to avoid imaging artifacts in perfusion and BOLD assessments, and iii) enabled finely tunable actuation for gradual, repeatable inflation and rapid recovery at different stages, addressing the limitations of conventional methods like middle cerebral artery occlusion (MCAO), photothrombosis, and clinical balloon catheters (**Fig. 1b–e; Extended Data Table 1**). These advancements make our device superior to other efforts in rodent hypoperfusion models, such as bilateral common carotid artery stenosis (BCAS) using microcoils^39^, which induce chronic ischemia but lack the temporal precision and reversibility needed for hyperacute studies. Clinical dual-lumen catheters, inflated at 6–8 atm, rely on folding of non-stretchable polymers, which constrains each model to one specific inflated diameter and results in limited elastic recovery^20,21^. Our micro-balloons show gradual, precise shape tunability and full reversibility within the required pressure. The single-shell design operates at lower pressures (1–3 atm), exerting a force up to ∼0.5 N to inflate up to ∼3 times its initial diameter. The double-shell design can exert a force ∼2.5 N, higher than a clinical catheter could do. Our micro-balloons also demonstrated effective intravascular occlusion in an in vitro model (700 μm outer diameter blocking 860 μm inner diameter lumens; **Extended Data Fig. 18, 19**), highlighting their potential for structural disruption in confined spaces without metallic interference in MRI. This design not only mitigates susceptibility artifacts common in BOLD and DSC sequences but also supports longitudinal imaging, a gap in traditional models where invasiveness precludes repeated assessments.

The ability to remotely modulate CCA occlusion allows us to monitor cerebral blood flow dynamics in real time using laser Doppler. Complete CCA occlusion induced a dramatic decrease in middle cerebral artery (MCA) flow (**Fig. 4c**), modeling ischemic stroke, cardiac arrest, hypoxic-ischemic encephalopathy, or severe systemic hypotension^40,41^. Notably, partial occlusions revealed a nonlinear, threshold-dependent response: significant flow reductions occurred only at ∼1/3 v constriction (**Fig. 4c,e**), challenging assumptions of proportional declines with blockage and suggesting compensatory mechanisms like autoregulation^42–44^. This nonlinearity may stem from collateral recruitment or myogenic adjustments in downstream arterioles, as observed in recent mouse models of partial reperfusion, where region-dependent pathophysiology delays neurodegeneration^45^. Post-occlusion hyperperfusion mirrored clinical observations in ∼50% of endovascular reperfusion patients (**Fig. 4c**)^46^, where it correlates with prolonged consciousness disturbances and hemorrhagic risk^47,48^, underscoring the need for models like ours to dissect these dynamics preclinically.

High-speed two-photon imaging further showed an 82% blood velocity reduction in large cortical blood vessels during complete bilateral occlusion of CCA, with residual flow (0.66 mm/s, **Fig. 5b**) attributed to vertebrobasilar contributions (20–30% of total cerebral supply), emphasizing regional vascular dependencies in mice, similar to humans^49^. This posterior circulation, involving vertebrobasilar arteries that supply the brainstem and cerebellar regions, exhibits parasympathetic regulation that may buffer hypoperfusion, as evidenced by human studies on posterior stroke hemodynamics^50^. Acute vasoconstriction observed in large cerebral arteries via TPFM aligns with rodent hypoperfusion models, potentially driven by endothelin-1 and prostaglandin EP1-mediated calcium influx in vascular smooth muscle^51,52^. This maladaptive response, including increased myogenic tone in parenchymal arterioles, underscores a shift toward constriction under reduced perfusion pressure, consistent with prior reports of velocity reductions during progressive carotid narrowing^53^. Our tunable occlusion platform allows dissection of these temporal sequences, revealing how initial autoregulatory failure could transition to compensatory dilation, which will inform therapies to target endothelin receptors.

In a multimodal MRI framework, we characterized hyperacute hemodynamic and metabolic responses to transient CCA occlusion. BOLD MRI revealed rapid, bilateral signal declines in multiple brain regions upon bilateral CCA blockage (**Fig. 6a–d**), aligned with previous findings that BOLD signal reductions reflect transient metabolic stress and increased deoxyhemoglobin during ischemic episodes^54,55^. DSC MRI mapped perfusion deficits via mean transit time (MTT), time-to-peak (TTP), and Tmax, confirming anterior-specific delays with little posterior involvement (**Fig. 6h**). The results are consistent with the anatomical distribution of blood supply (vertebrobasilar vs. internal carotid artery systems): anterior brain regions, including the frontal cortex and striatum, are predominantly supplied by the carotid circulation, while posterior structures such as the brainstem and cerebellum rely on vertebrobasilar arteries^56,57^. Our DSC MRI data uncovered asymmetric cerebral perfusion dynamics, with more pronounced delays in MTT and Tmax observed in the contralateral hemisphere compared to the ipsilateral side (**Fig. 6h**). This counter-intuitive pattern may arise from anatomical variations in the Circle of Willis or differential autoregulatory thresholds across hemispheres^58^. Together with perfusion asymmetries in clinical bilateral carotid stenosis where cross-hemispheric collaterals fail to fully compensate^59^, it underscores our model’s utility for probing variable vulnerabilities in hypoperfusion, which in turn will inform targeted therapeutic strategies to mitigate uneven risks in ischemic stroke.

Collectively, our remotely controlled elastomeric micro-balloons enable real-time in vivo imaging of the rodent brain’s hyperacute response to cerebral hypoperfusion across multiple modalities. The multimodal imaging platform consisting of through laser Doppler flowmetry, two-photon microscopy, BOLD MRI, and DSC MRI provides a reproducible framework for high-fidelity monitoring of blood flow, velocity, oxygenation, and perfusion dynamics during stage-wise, reversible ischemia. It surpasses single-modality studies by capturing spatiotemporal variations with unprecedented temporal precision at stroke onset. By enabling simultaneous occlusion and imaging, it sheds light on the “black box” of hyperacute stroke, revealing graded responses that could guide time-sensitive interventions. As the sample size of our pilot is small, *n* = 3–6 mice, larger cohorts for statistical power, and variability in Circle of Willis anatomy across strains should be screened via angiography in the future.

Beyond modeling hypoperfusion of the brain, our MRI-compatible micro-balloons hold translational potential for applications, including (i) modeling non-cerebral vascular stenoses (e.g., peripheral arterial, or aortic coarctation); (ii) sphincter dysfunctions (urinary, gastrointestinal, or pulmonary), and (iii) interventions like balloon kyphoplasty for vertebral restoration, urethral or esophageal dilation, or biliary obstruction relief. Recent applications of balloon guides in thrombectomy demonstrate improved outcomes in large-vessel occlusions, suggesting scalability to human neurovascular procedures. In non-neurological contexts, similar elastomeric designs could facilitate minimally invasive therapies for microvascular embolization or AVM embolization, where precise control minimizes off-target effects.

## Materials and Methods

### Materials

Sylgard 184 and Sylgard 186, consisting of hexamethyldisilazane (HMDS)-treated silica fillers, were purchased from Dow Chemicals. [(Mercaptopropyl)methylsiloxane]-dimethylsiloxane copolymer (SMS-142) was purchased from Gelest. 2-hydroxy-2-methylpropiophenone (Darocur 1173) was purchased from Sigma-Aldrich. Silica nanoparticles (diameter, 250 nm) are purchased from Sukgyung AT Co., Ltd. Ethanol (190 proof) was purchased from Thermo Fisher Scientific. Polyurethane acrylate (PU2100) was purchased from Miwon Chemicals. Blunt-tip needles (10 in. length) of gauge sizes 20G, 22G, and 24G were purchased from PATIKIL. All chemicals were used without further purification. Polyurethane external infusion tubing (VAHBPU-T25) was purchased from Instech Laboratories. Polyethylene micro-tubing (PE-8-100) was purchased from SAI. Norland Optical Adhesive NOA 68 was purchased from Norland Products. FITC-dextran (70 kDa; 90718) and agarose (A9539) were purchased from Sigma-Aldrich. Puralube was purchased from Dechra Veterinary Products. Circular coverslips (3 mm diameter) and donut-shaped coverslips (3.5 mm diameter) were purchased from Assistent Deckglaser. Isoflurane (Fluriso™, VetOne®, MWI Animal Health, Boise, ID, USA) and dexamethasone (2 mg/mL; VetOne®, MWI Animal Health, Boise, ID, USA) were obtained from VetOne; bupivacaine hydrochloride (0.25%, 2.5 mg/mL; Hospira, Inc., Lake Forest, IL, USA) was obtained from Hospira; and gadobutrol (Gadavist®, 1 mmol/mL; Bayer HealthCare Pharmaceuticals Inc., Whippany, NJ, USA) was obtained from Bayer.

### Fabrication of the PDMS microchannel mold

Two through-holes were drilled on the opposite sides of the side wall of a 140 mm-diameter plastic Petri dish (Thermo Fisher Scientific). Blunt-tip needles were positioned across the dish through the holes. Sylgard 184 prepolymer, mixed at a 5:1 weight ratio of base to curing agent, was then poured into the dish. A higher curing agent ratio than commonly reported in the literature was used to reduce the tackiness of the microchannel mold, facilitating the release of the elastomeric micro-balloons. After degassing in a vacuum chamber to remove air bubbles, the mixture was cured in a convection oven at 60 °C for 3 h. The Petri dish and needles were subsequently removed to obtain the PDMS mold.

### Formulation of silicone precursors

Sylgard 186, SMS-142, and 1 wt% of Darocur 1173 were mixed by a rotary mixer (ARE-310, Thinky) for 30 s. The molar ratio of the vinyl and thiol groups was set as 0.025, 0.05, 0.1, 0.2, and 0.4. For example, for 0.1 molar ratio, 1 g of Sylgard 186, 0.112 g of SMS-142, and 0.011 g of Darocur 1173 were mixed together.

### Formulation of PUA precursors

Silica nanoparticles were dispersed in ethanol (190 proof) for 3 h, followed by the addition of PU2100. The mixture was then sonicated for 10 min, and ethanol was evaporated in a convection oven at 60 °C for 12 h. As the loading of silica nanoparticles increased, the composite transitioned into a solid at a critical volume fraction, 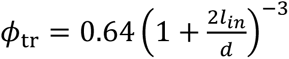^34^. The volume fraction of PU2100 was set to be 2 vol% lower than 𝜙_tr_ to ensure both a high *G*” and sufficient fluidity based on the previous literature^31^. After ethanol removal, ethanol and water were reintroduced into the system at 16 wt% and 4 wt%, respectively, relative to the mass of the silica nanoparticles, and mixed using the rotary mixer for 2 min.

### Fabrication of elastomeric micro-balloons

The silicone precursor was injected into the Sylgard 184 PDMS microchannel mold, followed by air injection to create a constant air front speed. For single-shell micro-balloons, the air front speed was set to 1 mm/s, regardless of the initial dimensions. Once the air core was formed, the precursor was UV-crosslinked by a UV-C LED strip light (270 nm, 1 mW/cm², Waveform Lighting) for 3 s to form the first shell. The short curing time was intended to leave unreacted functional groups that would bond with PUA precursor in preparation of the double-shell micro-balloons. In this case, the silicone shell was formed using the same method but at a reduced air front speed of 0.5 mm/s so that it was not too thick. After UV curing for 3 s, the PUA precursor was injected into the air core, leaving a defined length unfilled near the tip of the tube. Air was injected again at 1 mm/s, followed by UV curing, and the same procedure was repeated at the opposite end of the tube to define the length of the balloon segment. Finally, the double-shell tube was UV cured for another 10 min under nitrogen, after which the PDMS mold was removed by cutting. The released micro-balloon was connected to a blunt tip nozzle using a UV-curable adhesive (Photobond GB368, Delo). Although the microchannel with an outer diameter as small as 550 μm could be fabricated, its small inner diameter (∼200 μm) made it difficult to accommodate standard needles. Nevertheless, the high elasticity of the PUA shell allows a needle to be inserted into the microchannel to function as a rigid nozzle (**Extended Data Fig. 20**).

### Characterizations

The micro-balloons were imaged under the optical microscope (BX61, Olympus) in the transmission mode. For the longitudinal views, micro-balloons were embedded in the mold, which had the same refractive index, thus minimizing interfacial diffraction for direct observation of the wall thickness. The microstructures of the micro-balloons with silica nanofillers/nanoparticles were characterized by the high-resolution scanning electron microscopy (7500F HRSEM, JEOL).

### Assembling a micro-balloon with a syringe

Polyurethane external infusion tubing (∼ 200 cm long) and polyethylene micro-tubing (∼ 4 cm long) were prepared. A 1.0 cm segment of PE-8-100 was inserted into the T25 tubing, and NOA 68 was applied at the junction and cured under 365–405 nm UV illumination for 1 min. Subsequently, 0.5 cm of PE-8-100 was inserted into the double shell, and the joint was sealed with hot melt adhesives (3M, St. Paul, MN, USA). The assembly was then filled with water using a syringe until the tip end was saturated, followed by hot glue sealing. A plastic C-ring, cut from a plastic tube, was positioned around the micro-balloon.

### Mice

All experimental protocols at the University of Maryland, Baltimore were conducted according to the National Institutes of Health (NIH) guidelines for animal research and approved by the Institutional Animal Care and Use Committee at the University of Maryland, Baltimore. Mice (3-4 months old, 25-30 g, VR breeding) were group housed with littermates until craniotomy surgery, after which they were singly housed. Mice were maintained on a 12– 12-h (6 a.m.–6 p.m.) light–dark cycle. They were anesthetized with isoflurane (4-5% for induction, 1.5% for maintenance).

### Surgical procedure for micro-balloon occlusion and cerebral blood flow monitoring by laser Doppler flowmetry

After anesthetizing, mice were placed in a lateral position on the homoeothermic-heating pad (Physio Suite Mouse STAT Right Temp, Kent Scientific Co., Torrington, CT), and the Puralube was applied to the eyes. A 5 mm scalp incision between the right eye and ear was made to expose the temporal bone to insert a needle laser Doppler probe (diameter, 2 mm) (Moor Instruments, Inc., Wilmington, DE). The temporalis muscle was cauterized to visualize the middle cerebral artery (MCA). Mice were then placed in the supine position, where the neck and chest were disinfected. A 2-cm midline neck incision was made to expose both the common carotid arteries (CCAs), avoiding injury to the vagus nerve. The left CCA was permanently ligated using a 7-0 monofilament suture, and the micro-balloon connected to a Hamilton syringe was placed around the right CCA (see **Fig. 4a**). A laser Doppler probe was positioned over the left MCA territory to monitor the cerebral blood flow. Stepwise inflation of the micro-balloon with water was performed to achieve progressive attenuation of the blood flow. Doppler recordings were collected continuously from the baseline through occlusion and reperfusion, with at least 2 min between occlusions. After the procedure, the Doppler probe was removed, wounds were sutured, and mice were recovered on a 37°C heating pad before returning to their cages. For two-photon imaging and MRI, the exposed neck region was filled with 2% agarose to stabilize the micro-balloon, followed by suturing the skin to further stabilize the catheters. For mice with cranial window, the Doppler probe was mounted on a holder and gently positioned to contact the cranial window.

### Cranial window surgery

Mice were anesthetized and positioned in a stereotaxic frame (RWD, Sugar Land, TX) equipped with a homeothermic heating system. The scalp was treated with 70% ethanol and given a subcutaneous injection of 2.5 mg/mL bupivacaine and 2 mg/kg dexamethasone prior to the surgery. A circular craniotomy (3 mm diameter) was performed over the left primary visual cortex (V1). Afterward, a 3-mm diameter circular coverslip adhered to a 3.5-mm diameter donut-shaped coverslip affixed over the craniotomy using NOA 68, and a custom titanium headpost was then secured to the skull with dental cement. Mice were allowed to recover on a heating pad before being returned to their home cages. Mice received subcutaneous injections of carprofen (5 mg/kg, Rimadyl, Zoetis) pre-operatively and once daily for three consecutive days. Laser Doppler flowmetry or two-photon imaging was performed at least 14 days after cranial window implantation to minimize the effects of acute postoperative inflammation on vasomotion.

### Two-photon line-scan and imaging

Mice implanted with a micro-balloon catheter were anesthetized and placed on a heating blanket to maintain body temperature at 37 °C. To visualize the vasculature, 1.0 % FITC-dextran was administered via retro-orbital sinus injection. TPFM was performed using a custom-built two-photon microscope using a water dipping objective (XL Plan N 25x/1.05 W MP 0-0.23 OFN18 Olympus). Two-photon excitation was obtained using an 80 MHz Ti-Sapphire laser (Chameleon Discovery, Coherent) with sufficient power from 710 nm to 1,080 nm. The laser was tuned to an excitation wavelength of 940 nm to optimize the fluorescence excitation of FITC-dextran-labeled vessels. For blood flow velocity measurements, line-scan imaging was performed along the central axis of target vessels at a sampling rate of 1 kHz. In the resulting space–time images, the slope of red blood cell (RBC) streaks reflected flow velocity. The orientation and length of the scan line were manually adjusted to align with the vessel centerline. Data acquisition was controlled using ScanImage software (Vidrio Technologies), and subsequent blood flow velocity analysis was performed using custom-written scripts in MATLAB^60^.

### MRI Acquisition and Perfusion Mapping

Mice with the micro-balloon secured around the left CCA were positioned in a 9.4 T MRI scanner (Bruker). Baseline T2-weighted (TR/TE = 2500/10 ms) and T1-weighted (TR/TE = 1500/7.5 ms) images were acquired. Stepwise inflation and deflation of the micro-balloon were performed to transiently occlude the CCA during MRI acquisition. Functional MRI data for detecting oxygenation changes were acquired using a gradient-echo echo-planar imaging (GE-EPI) sequence (GEEPI_dyn_cor, based on EPI.ppg pulse program) on a Bruker BioSpec preclinical MRI system operating at 9.4 T. Images were obtained in the axial plane with a dynamic time-series acquisition (3 repetitions) to capture BOLD contrast. Key sequence parameters included a repetition time (TR) of 500 ms, echo time (TE) of 11.76 ms, number of averages of 1, field of view (FOV) of 15.9 × 15 mm, acquisition matrix size of 104 × 128 (reconstructed to 128 × 128), 11 slices with a thickness of 0.7 mm and inter-slice separation of 0.95 mm, in-plane resolution of 0.124 × 0.117 mm, and a total data depth of 33 (reflecting 11 slices across 3 dynamic repetitions). The data were stored as 16-bit signed integers in little-endian byte order. Data were preprocessed to correct for motion and distortions prior to analysis of perfusion dynamics and BOLD signal changes. A programmable syringe pump (PHD 2000, Harvard Apparatus) was used to deliver gadolinium contrast agent (Gadavist®, diluted 1:10 in saline) via the tail vein at 0.6 ml/min during dynamic T1-weighted imaging (TR/TE = 200/5.2 ms; field of view, 15 × 15 mm²; matrix, 128 × 64; 50 repetitions; acquisition time, 240 s). A total of 70 µl was infused immediately after balloon inflation to visualize the perfusion territory. The same GE-EPI sequence as above was used for acquiring the signal changes post-gadolinium injection in DSC perfusion MRI.

### MRI data processing

Time-series MRI for oxygenation and Gd-dynamics analyses were processed in ImageJ (v1.54, NIH). The baseline and subsequent occlusion epochs were defined as in the schematic timeline Regions-of-interst (ROIs) (cortex, hippocampus, striatum, and vertebrobasilar territory) were manually placed in predefined anatomical regions, and mean signal intensities were extracted over time using the *Stack Plot Z-axis* function. For each ROI, values corresponding to the micro-balloon inflation and deflation phases (6–30 s after onset) were averaged and normalized to the corresponding baseline mean (pre-inflation) to yield normalized oxygenation traces. Normalized time courses were plotted in GraphPad Prism (v10.3.0, GraphPad Software) to generate dynamic oxygenation curves. Perfusion parameters were analyzed using Horos (v3.3.6; The Horos Project, USA), an open-source DICOM viewer. Dynamic susceptibility contrast (DSC) MRI datasets were imported into Horos and processed with the IB Neuro plugin. An arterial input function (AIF) was manually placed in a small cortical artery. Parametric maps—relative cerebral blood flow (rCBF; from the deconvolved residue function), mean transit time (MTT), and time to maximum (Tmax; delay to the peak of the deconvolved residue)—were generated using IB Neuro’s standard deconvolution pipeline, with leakage correction enabled. Bilateral target ROIs (cortex, hippocampus, striatum, and vertebrobasilar territory) were manually delineated in Horos on representative slices and applied consistently across conditions. Mean parameter values were extracted per ROI and analyzed in GraphPad Prism the Open–Blocked difference was summarized as reduction % = (Open − Blocked)/Open × 100%.

### Statistical analysis

Data analysis was performed using MATLAB (MathWorks), Prism (v10.3.0, GraphPad Software) and PAST (v4.14). Normality was assessed using the Shapiro– Wilk test. For normally distributed paired data (Fig. 5g; Fig. 6d, except striatum; and Fig. 6g), parametric paired t-tests were used. For non-normally distributed paired data (Fig. 4d, e,h; Fig. 5c; and striatum in Fig. 6d), Wilcoxon matched-pairs signed-rank tests were applied. Differences in Fig. 6h–j were analyzed using one-sample t-tests. Normally distributed data are presented as mean ± standard deviation, in bar plots, whereas non-normally distributed data are presented as median ± IQR in boxplots (median, 25th–75th percentiles; whiskers, Tukey style ± 1.5× IQR). Statistical significance was defined as *\* p <* 0.05, *\*\* p <* 0.01, *\*\*\* p <* 0.001, and **** p < 0.0001. Medians, IQRs, means and s.e.m. values are reported throughout the text.

## Acknowledgements

The research is supported in part by the National Institutes of Health (R21AG077631; R03NS123733; R03NS128459; R21AG074978 to Y.L. and R01DA056739 to P.W.), Maryland Stem Cell Research Fund (2024-MSCRFD-6363 and 2022-MSCRFL-5893 to Y.L. and 2022-MSCRFD-5886 to P.W.), and National Science Foundation (NSF)’s Materials Research Science and Engineering Center (MRSEC) at University of Pennsylvania (DMR-2309043, to S.Y.). Jinghui W. acknowledges postdoc fellowship from Maryland Stem Cell Research Fund (2024-MSCRFF-6328). The Center for Innovative Biomedical Resources (CIBR) at the University of Maryland School of Medicine is acknowledged for access to imaging instrumentation and expertise. Illustrative elements in this manuscript were created using academic resources from BioRender (www.biorender.com).

## Author contributions

S.Y., Y.L and P.W. conceived the idea and designed the research. J.B.K. and Jinghui.W. contributed equally to this work. J.B.K. devised the micro-balloon design and material formulations, fabricated the micro-balloons, and characterized their dimensional change and expansion forces. Jinghui.W. performed in vivo experiments and analysis with the help of H. T, and G.Q. under the supervision of Y.L, P.W. and M. J. P.W. performed and supervised MRI experiments and analysis. Y.C. characterized the mechanical properties of micro-balloons. Jingxian.W. performed the adhesion test. A.N. tested the rheological properties of the micro-balloon materials. J.B.K., Jinghui.W., Y.L., and S.Y. wrote the manuscript. All authors participated in discussions and reviewed the manuscript.

## Additional information

Extended data is available for this paper online.

Supplementary information is available in the online version of the paper.

Correspondence and requests for materials should be addressed to P.W., Y.L., and S.Y.

Reprints and permissions information is available online at www.nature.com/reprints.

---

**Extended Data Fig. 1.**
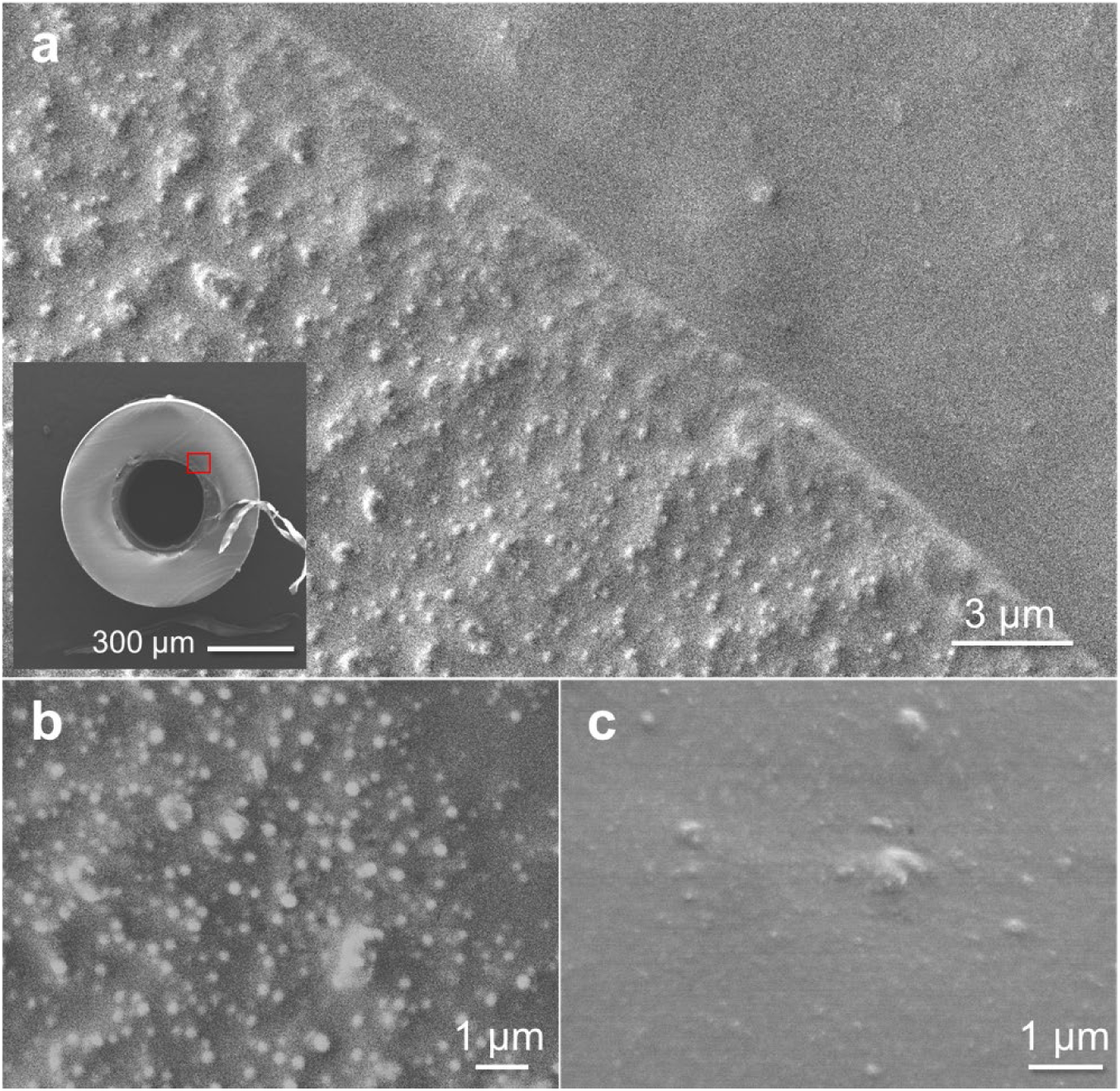
Silica fillers/nanoparticles inside the micro-balloons. **a–c,** Cross-sectional SEM images at the interface between the silicone and PUA shells (a), with magnified views of the PUA (b) and silicone (c) shells. The inset in (a) shows a low-magnification cross-section image with the red box indicating the region corresponding to a.

**Extended Data Fig. 2.**
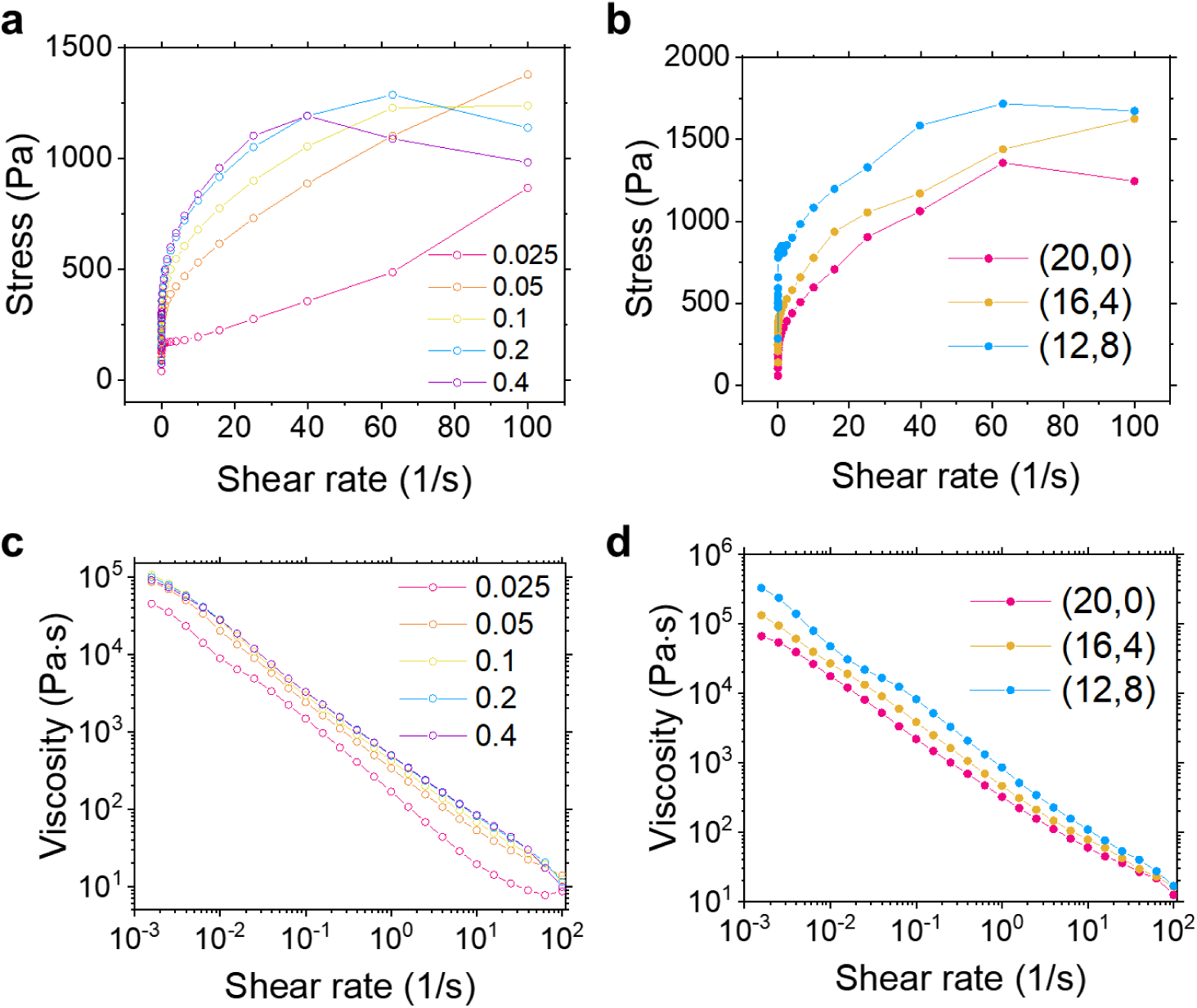
Rheological properties of the precursors. **a,b,** Shear rate-dependent yield stress of (a) silicone and (b) PUA precursors with different compositions. **c,d,** Shear rate-dependent viscosities of the precursors for (c) silicone and (d) PUA with different compositions.

**Extended Data Fig. 3.**
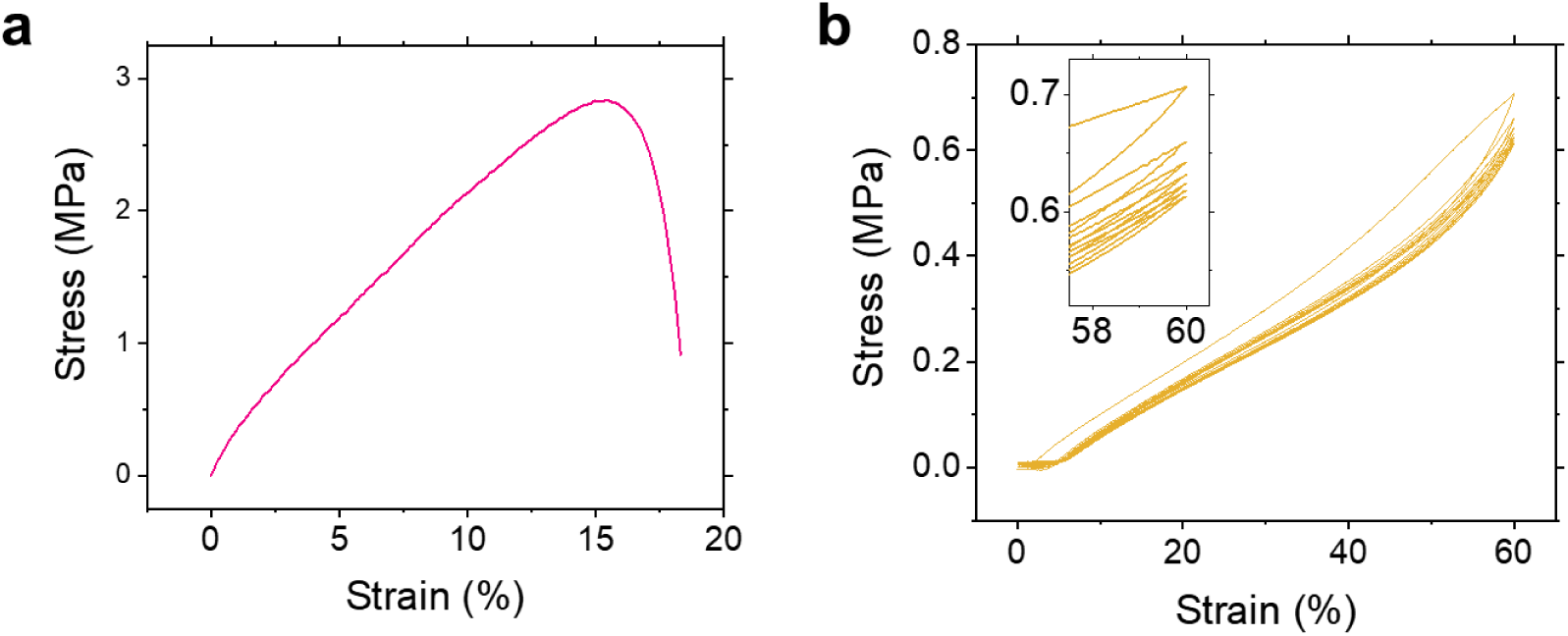
Tensile tests of the shell materials. **a,** Pure crosslinked PUA without silica nanoparticle. **b,** Cyclic tensile test of the silicone precursor with m_=_/m_SH_ of 0.1 at 60% strain.

**Extended Data Fig. 4.**
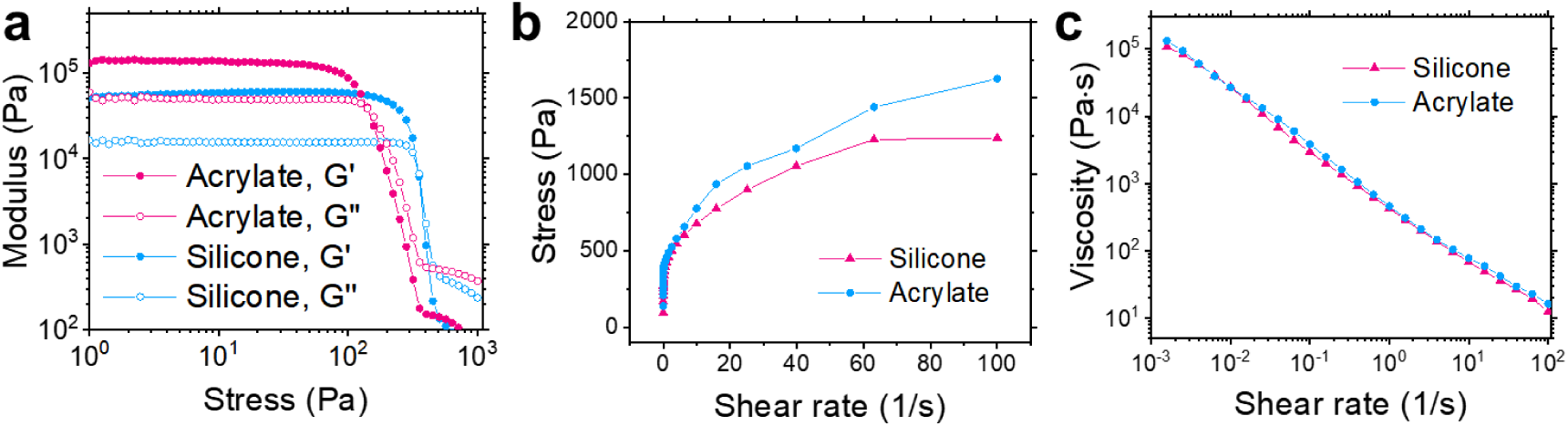
Rheological properties of the optimal precursors. **a,** Oscillatory shear moduli. **b,** Shear stress versus shear rate. **c,** Shear-rate-dependent viscosity.

**Extended Data Fig. 5.**
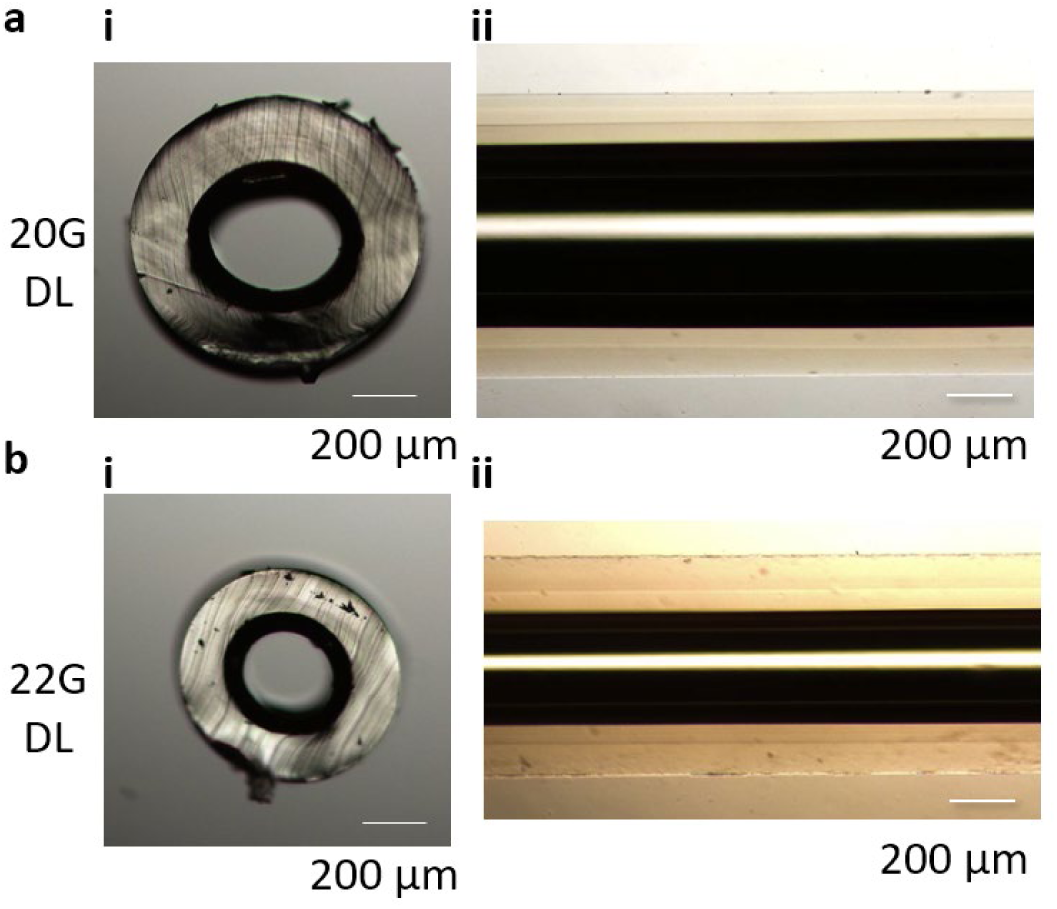
Optical microscopy (OM) images of the double-shell micro-balloons. **a,** Cross-sectional view. **b,** Top view.

**Extended Data Fig. 6.**
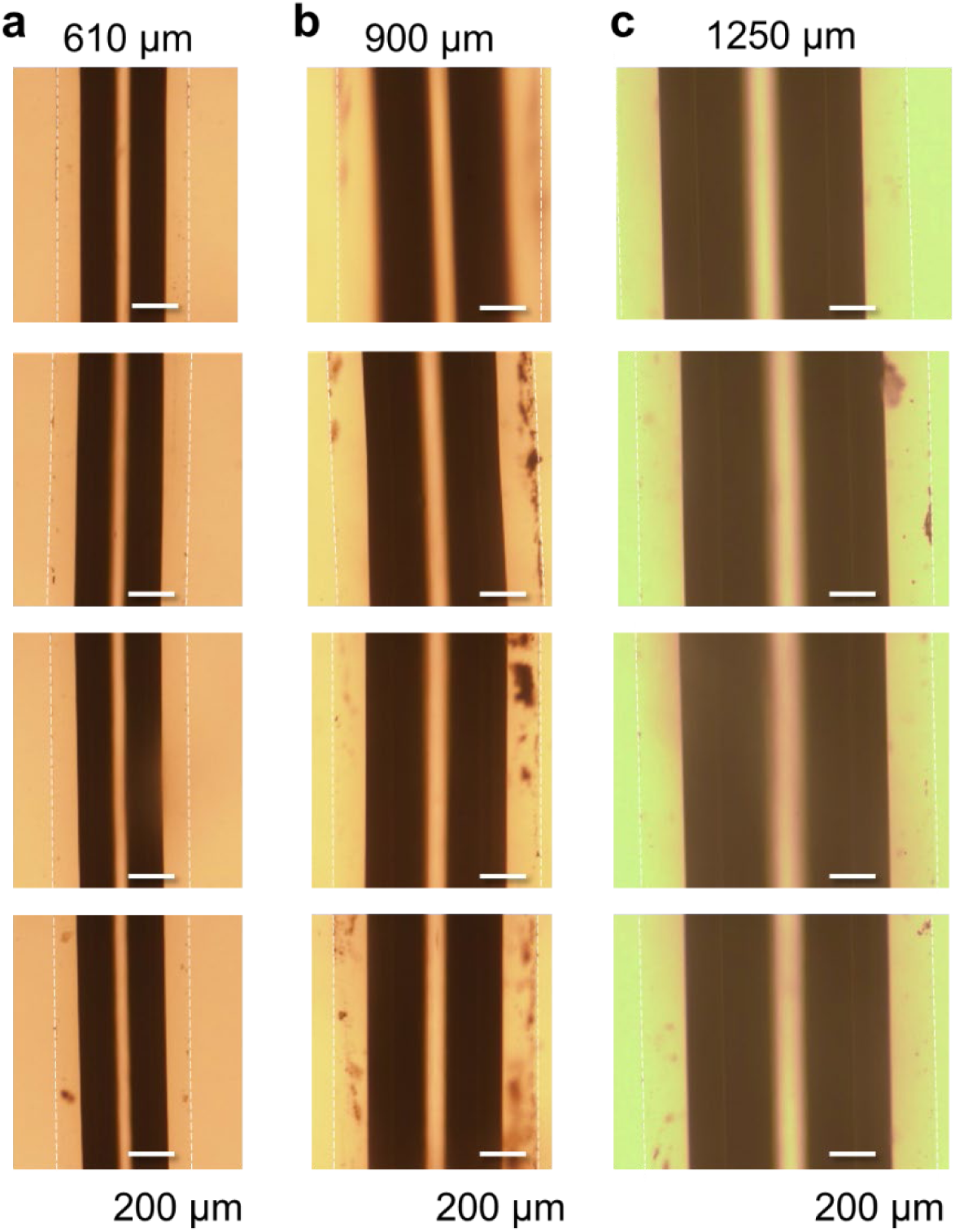
OM images of the micro-balloons fabricated by a constant air injection speed of 1 mm/s for varying channel inner diameters, demonstrating structural uniformity. **a,** 610 μm, **b,** 900 μm, and **c,** 1250 μm.

**Extended Data Fig. 7.**
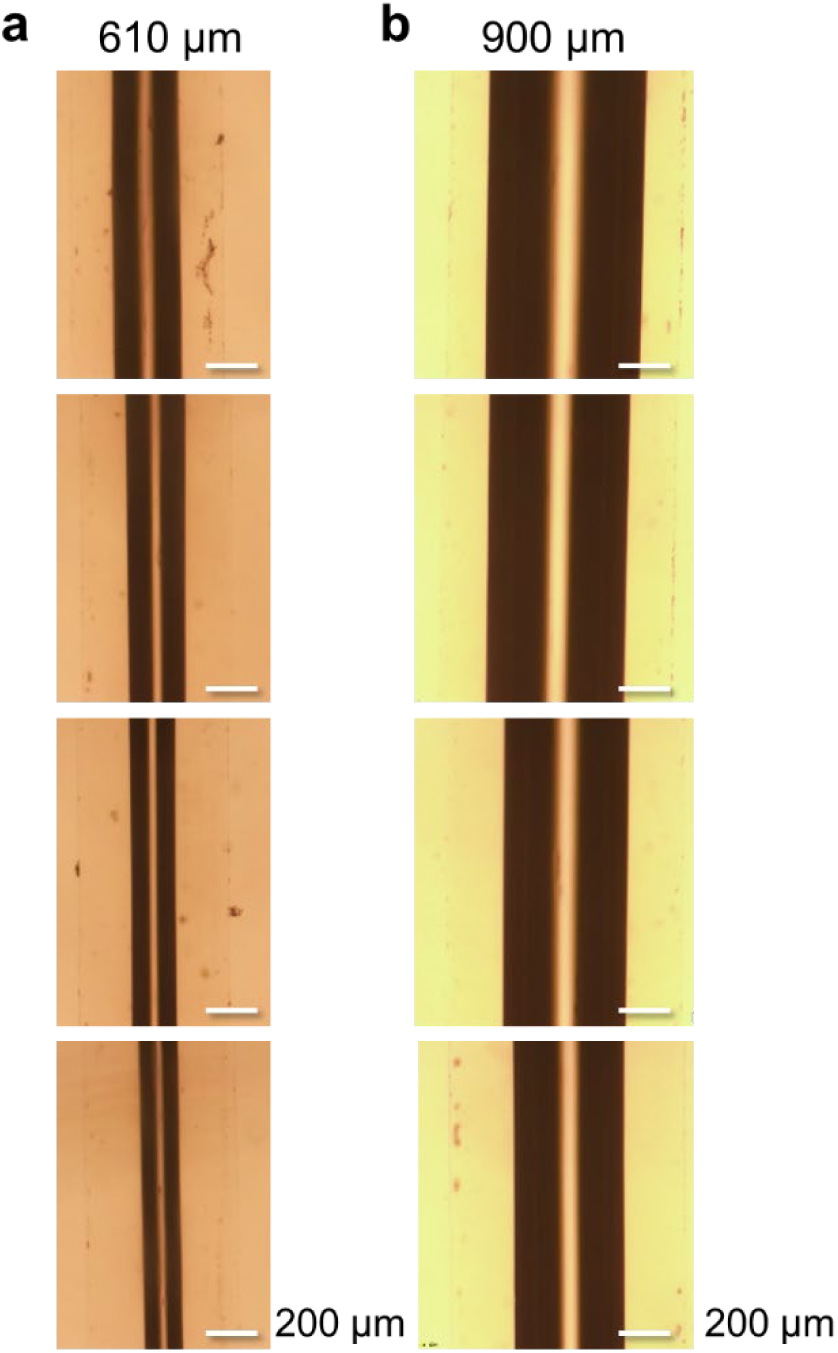
OM images of the micro-balloons fabricated by a constant air pressure of 1.3 mm, demonstrating structural non-uniformity. **a,** 610 μm, **b,** 900 μm.

**Extended Data Fig. 8.**
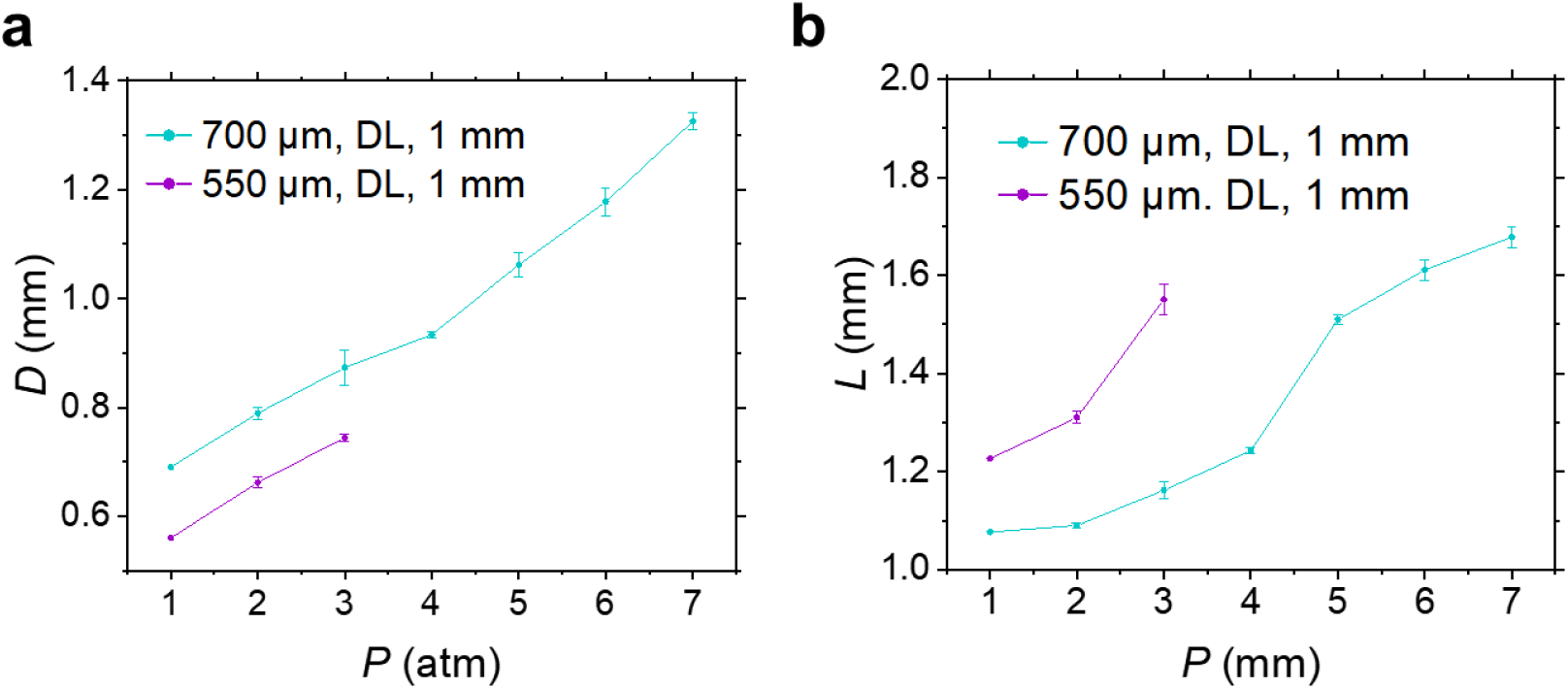
Pressure-dependent dimensional changes of the double-shell micro-balloons. **a,** Changes in the diameter (*n* = 3). **b,** Change in length. The micro-balloons have outer diameters of 700 μm and 550 μm, respectively (*n* = 3).

**Extended Data Fig. 9.**
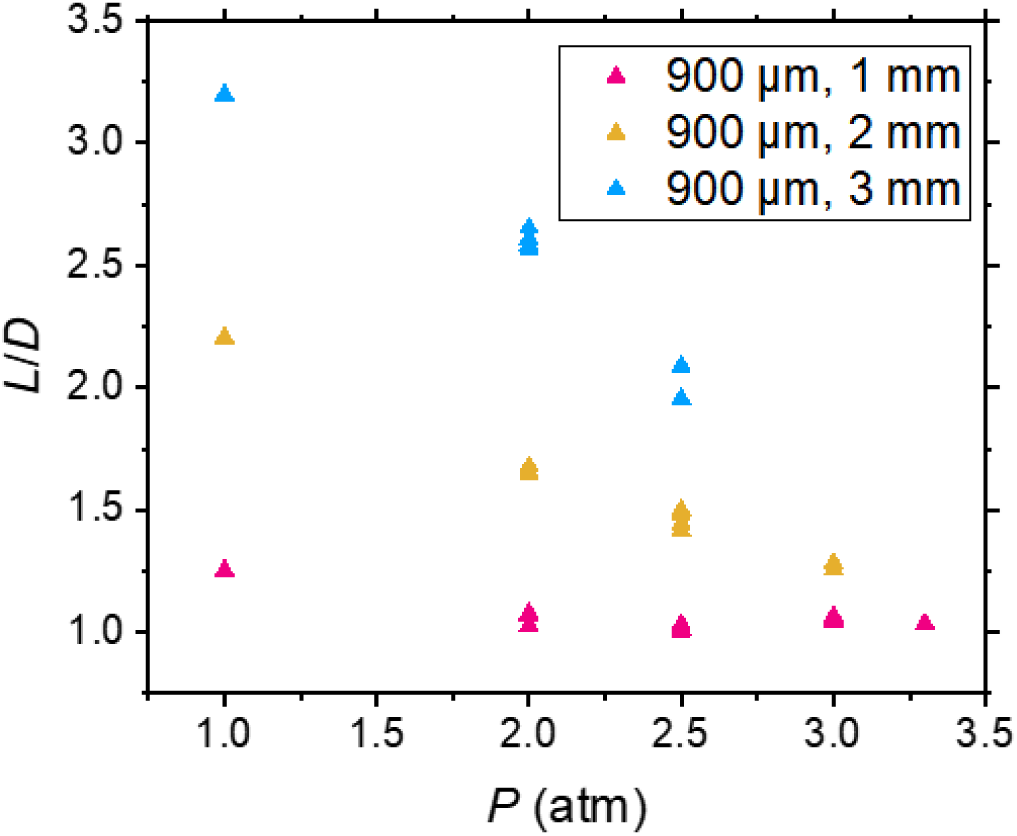
Pressure-dependent aspect ratios (length-to-diameter, L/D) of inflating micro-balloons with varying balloon segment lengths.

**Extended Data Fig. 10.**
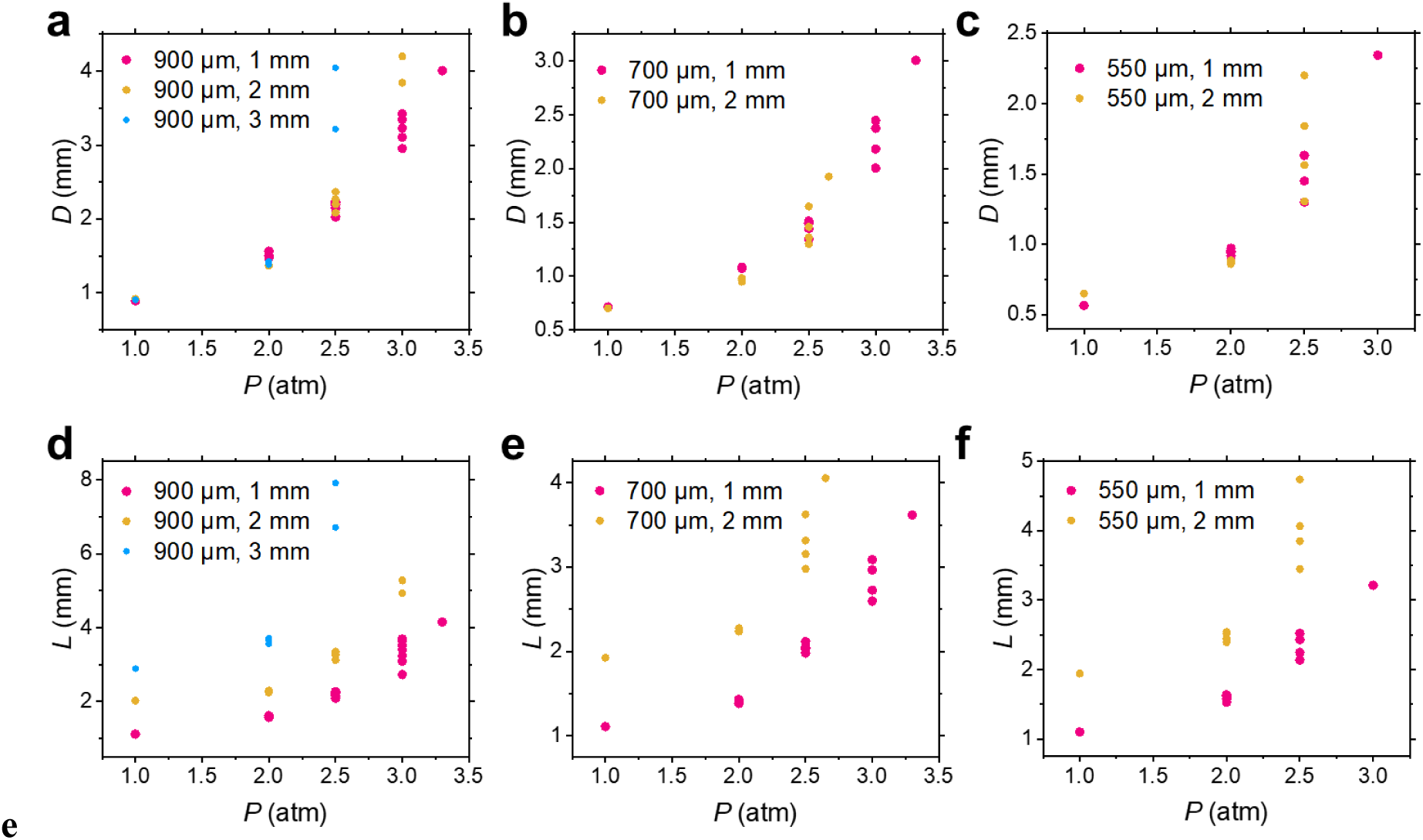
Pressure-dependent dimensional changes of micro-balloons with varying length. **a–c,** Changes in the diameter. **d–f,** Change in the length. The balloon segments with varying outer diameters (900, 700, 500 μm) and balloon segment lengths (1, 2, 3 mm) (n = 3).

**Extended Data Fig. 11.**
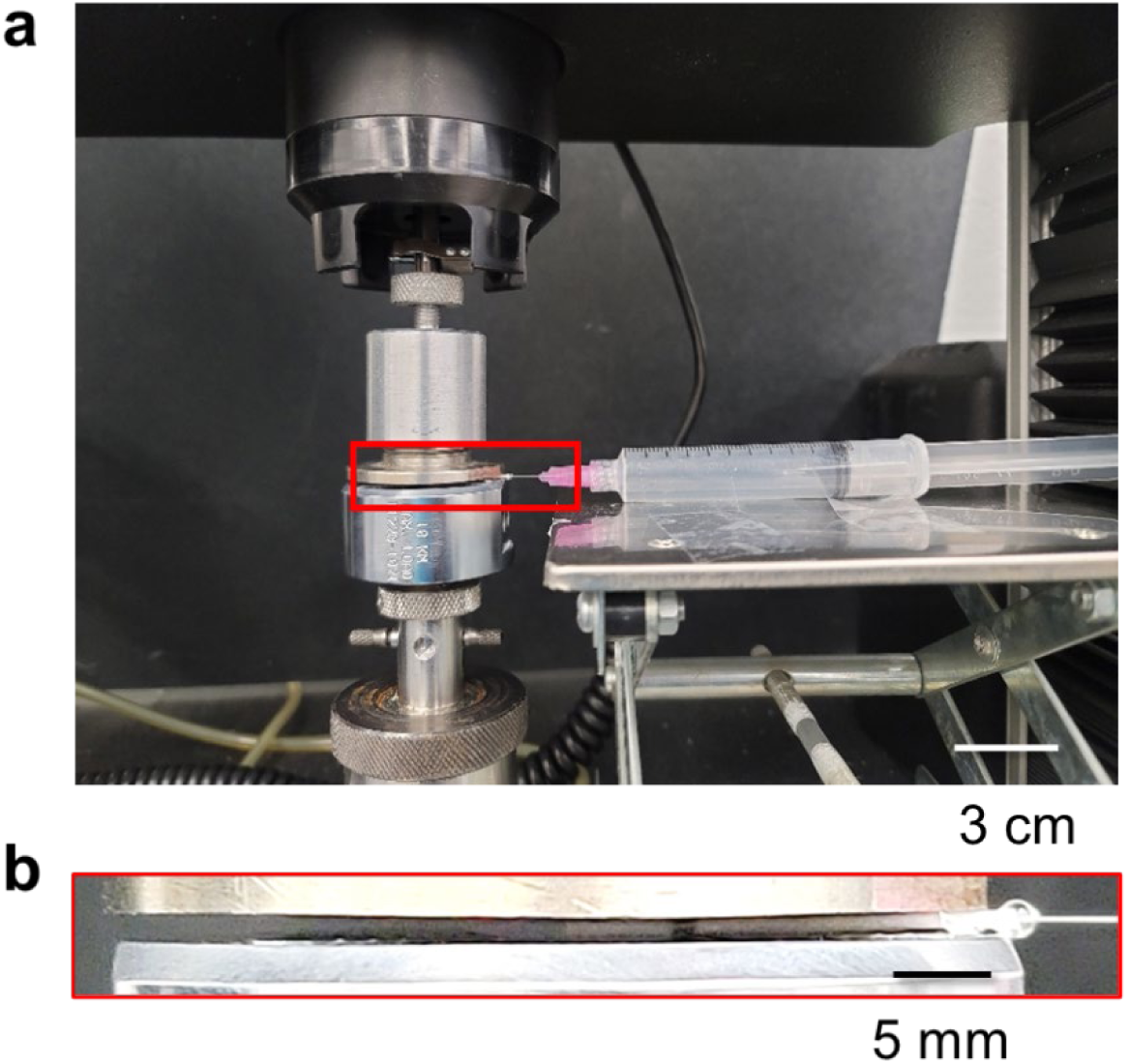
Setup to measure the expansion force of micro-balloons using a universal testing machine. **a,** Full view. **b,** close-up view.

**Extended Data Fig. 12.**
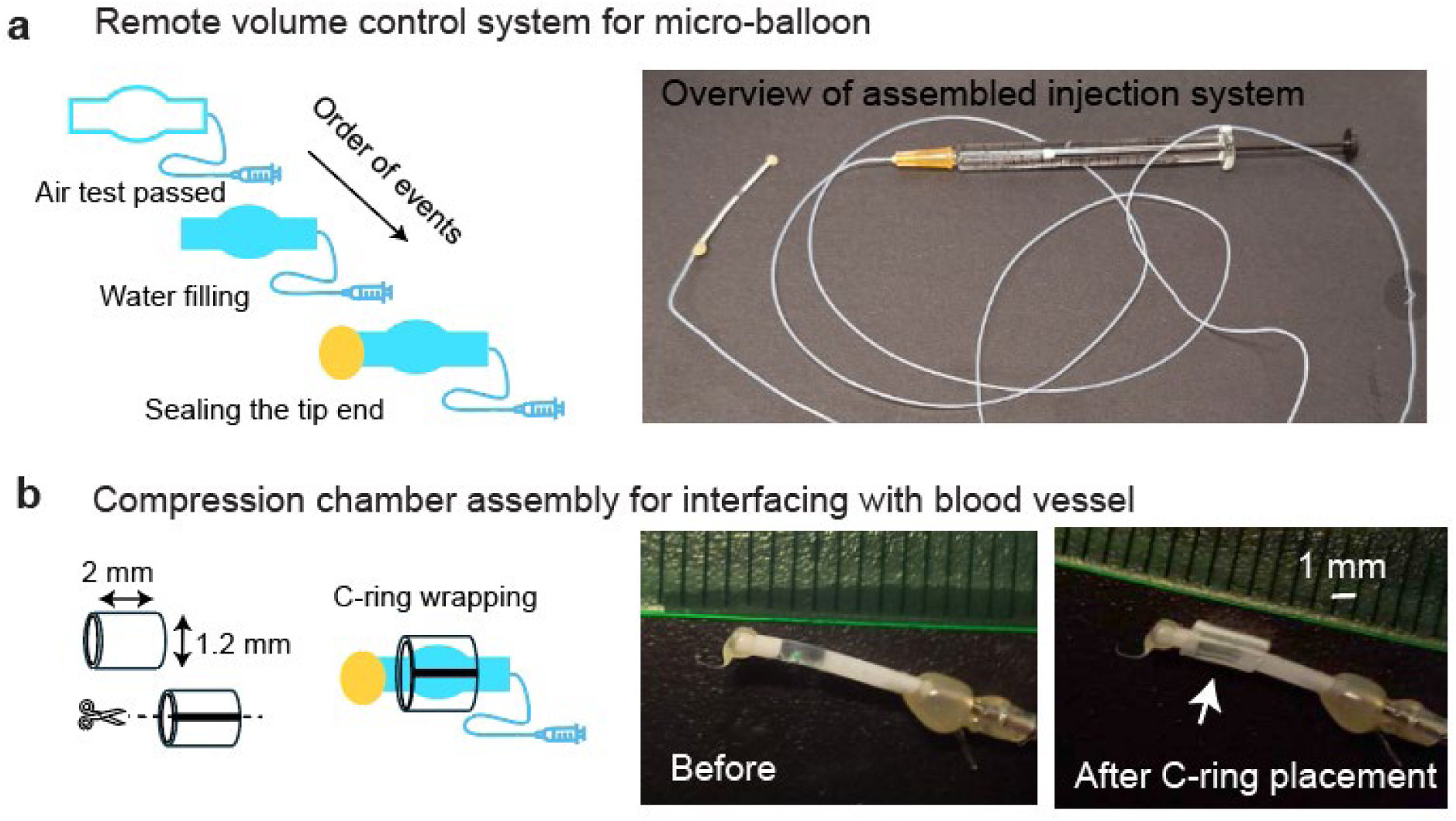
Assembly of the remote micro-balloon control system. **a,** Hydraulic micro-balloon volume control. Micro-balloons that passed the air test were connected to a Hamilton syringe through a long tube and filled with water. The tip end was sealed with hot glue. **b**, Compression chamber assembly. A C ring was made from a plastic tube to surround the micro-balloon.

**Extended Data Fig. 13.**
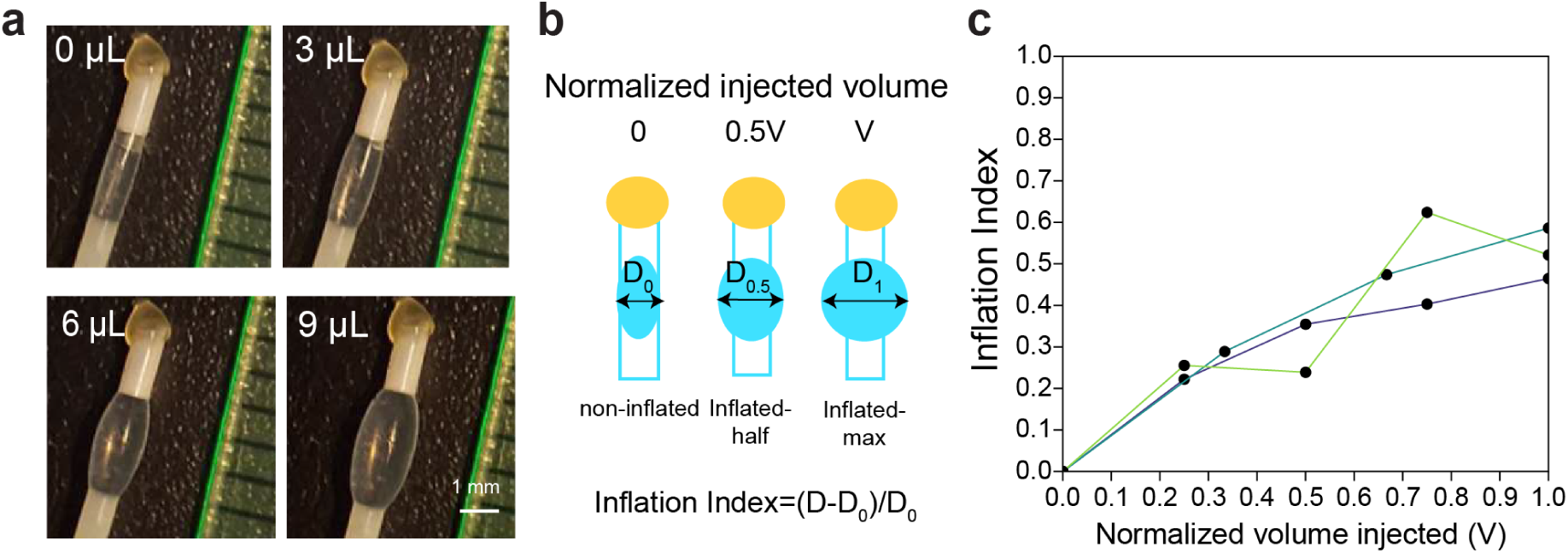
Expansion of the micro-balloon vs. the volume of water injected. **a**, A representative micro-ballon inflated by different amounts of water. **b**, Schematic illustration of how Inflation Index is calculated: the ratio of increased diameter to the starting diameter. V represents the maximum volume that is injected for inflation of the micro-ballon. **c**, Inflation Index as a function of the normalized injection volume (n = 3).

**Extended Data Fig. 14.**
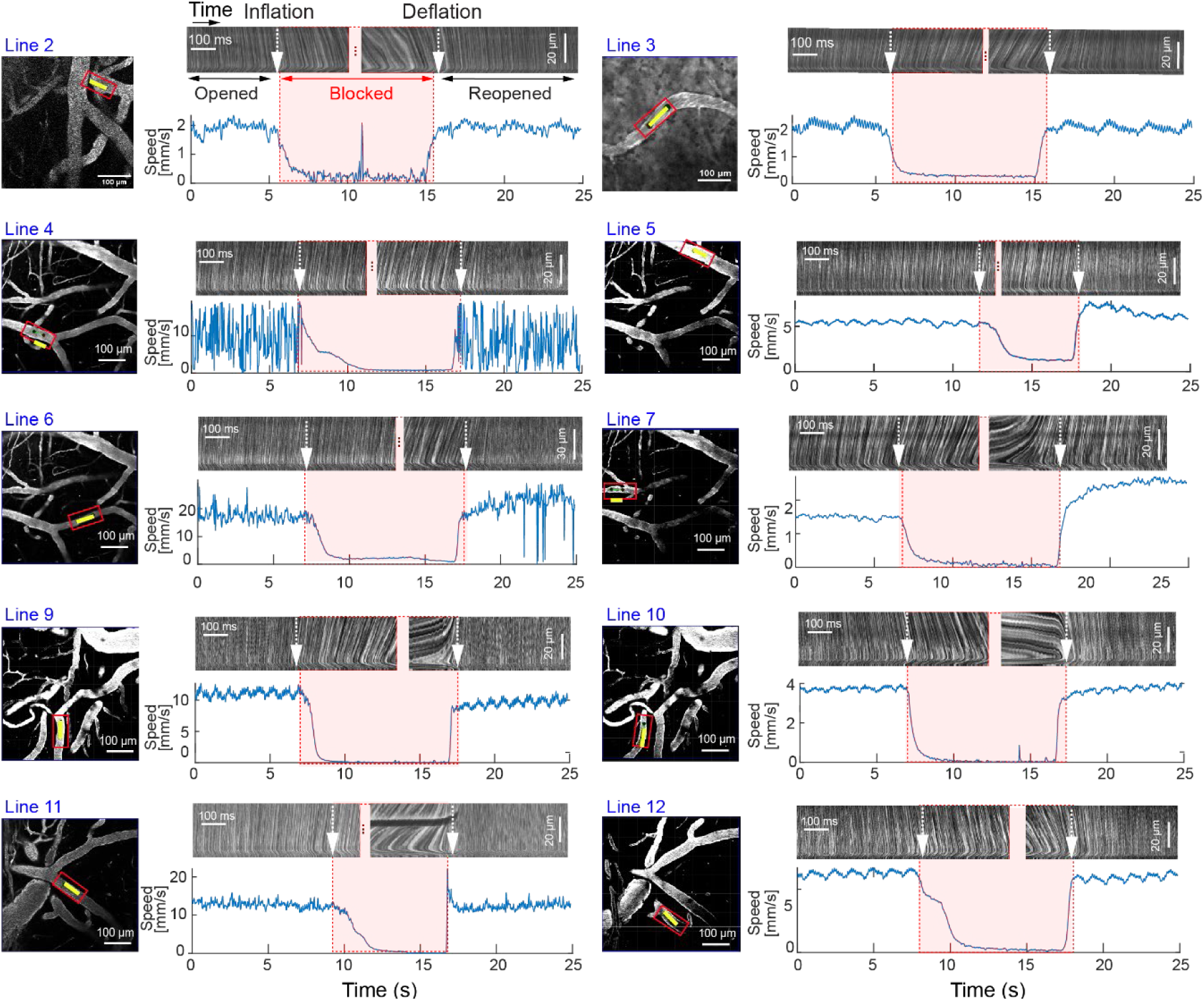
More examples of the line-scans over the blood vessels. Two-photon line-scan images (top panels) and corresponding blood flow velocity traces (bottom panels, blue color) from vessels in multiple field-of-view (shown on the left) mice before, during, and after micro-balloon occlusion of the common carotid artery (CCA). Measurement sites are indicated by red boxes on vascular images (left panels). The timing of inflation and deflation is indicated above each trace. Lines 2–3 correspond to recordings from mouse 1, lines 4–7 from mouse 2, and lines 9–12 from mouse 3. Scale bars, 100 μm (vascular images) and 100 ms (traces panels).

**Extended Data Fig. 15.**
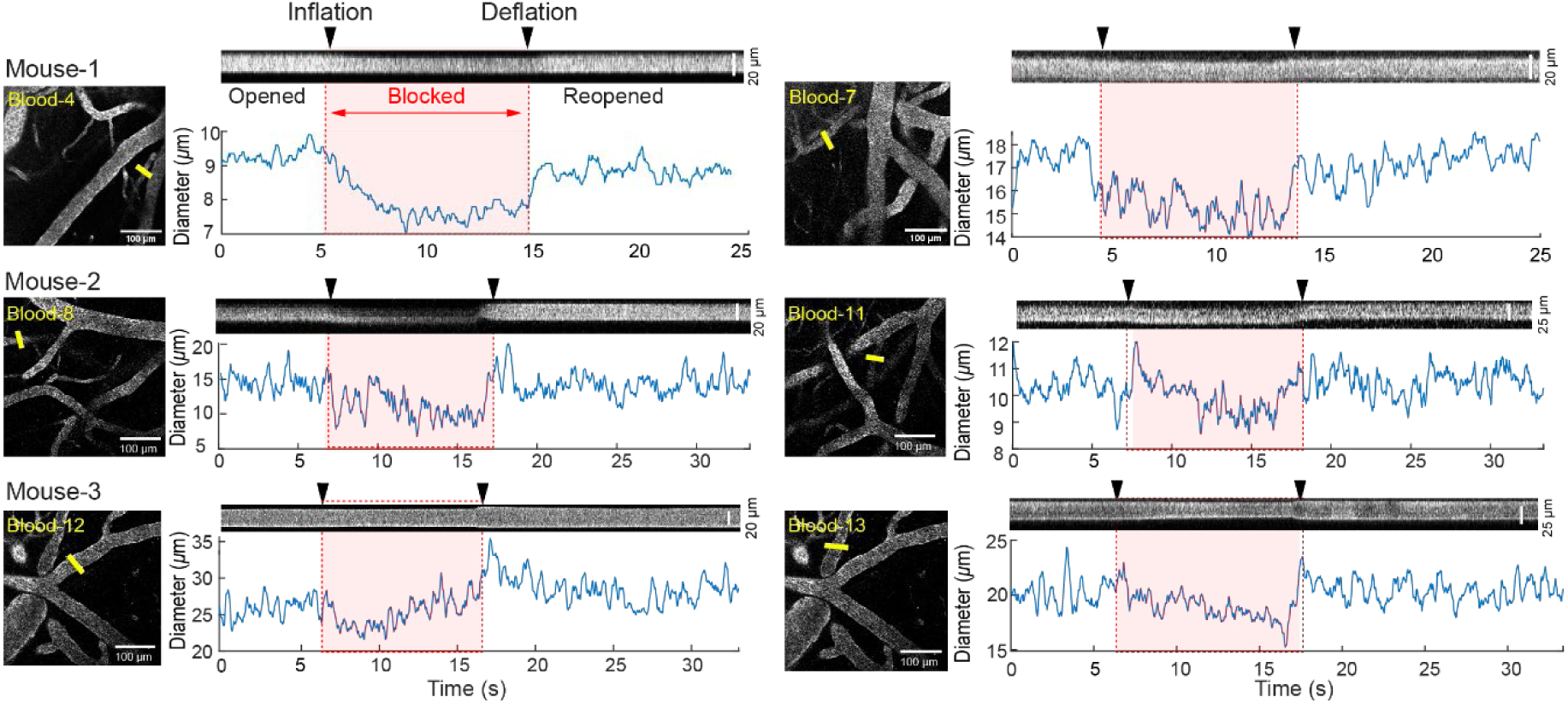
Blood vessel diameter changes during micro-balloon–induced CCA occlusion. Representative two-photon images of cortical vessels (left) with yellow arrows indicating measurement sites, and corresponding diameter traces (right) before, during, and after micro-balloon occlusion of the common carotid artery (CCA). Time points for non-blocked, inflation, blocked, deflation, and reopened states are indicated above each trace. Scale bars, 100 μm.

**Extended Data Fig. 16.**
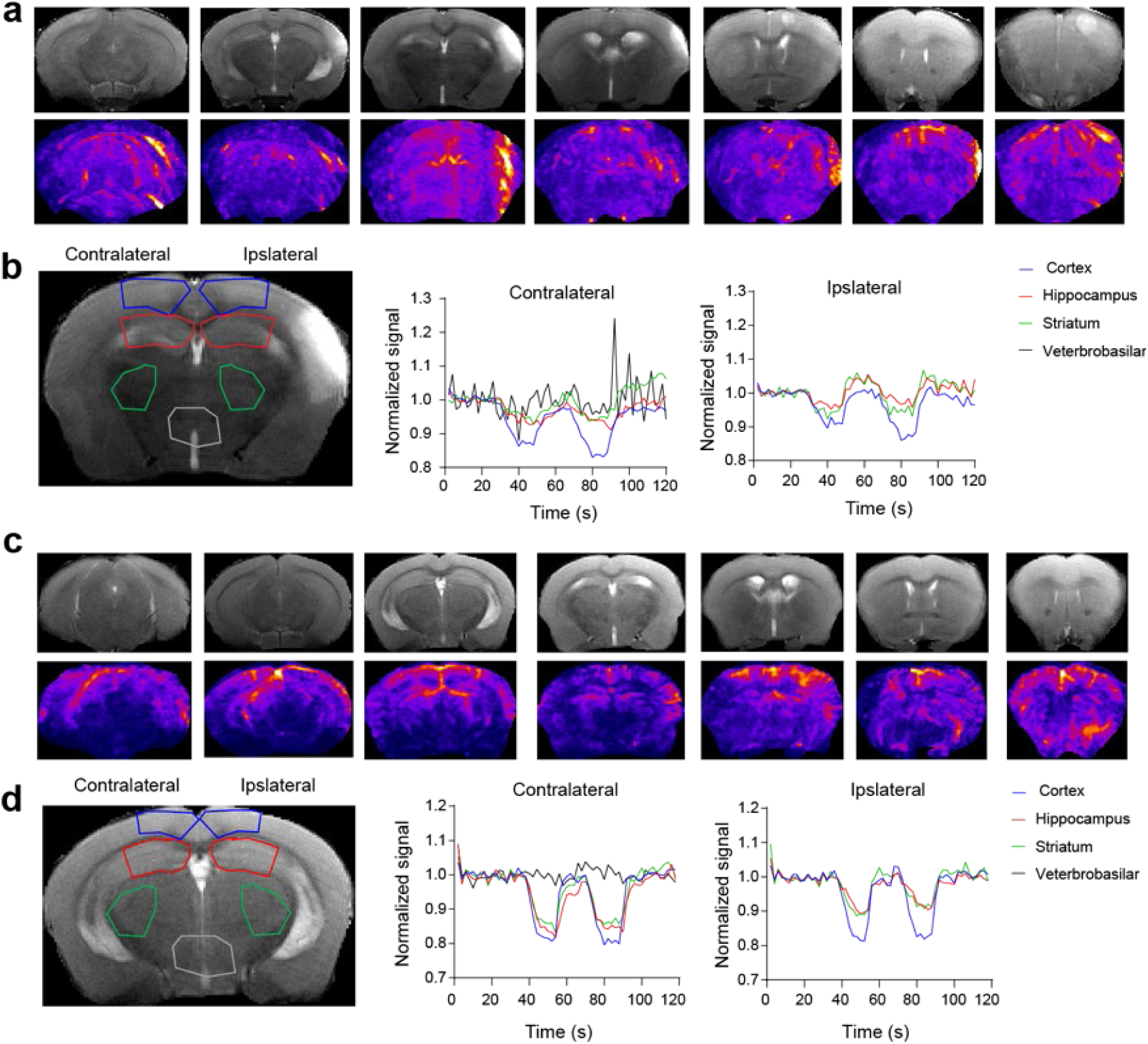
Additional examples of oxygenation dynamics during micro-balloon–induced CCA occlusion. **a,c,** Representative coronal T2-weighted images (top) and corresponding oxygenation standard deviation (s.t.d.) maps (bottom) from two additional mice during CCA occlusion. **b,d,** Example coronal sections showing the regions of interest (ROIs; cortex, hippocampus, striatum, and vertebrobasilar territory) outlined on T2 images (left) and the corresponding normalized oxygenation time courses for each ROI (right). Warmer colors in the s.t.d. maps indicate higher signal variability.

**Extended Data Fig. 17:**
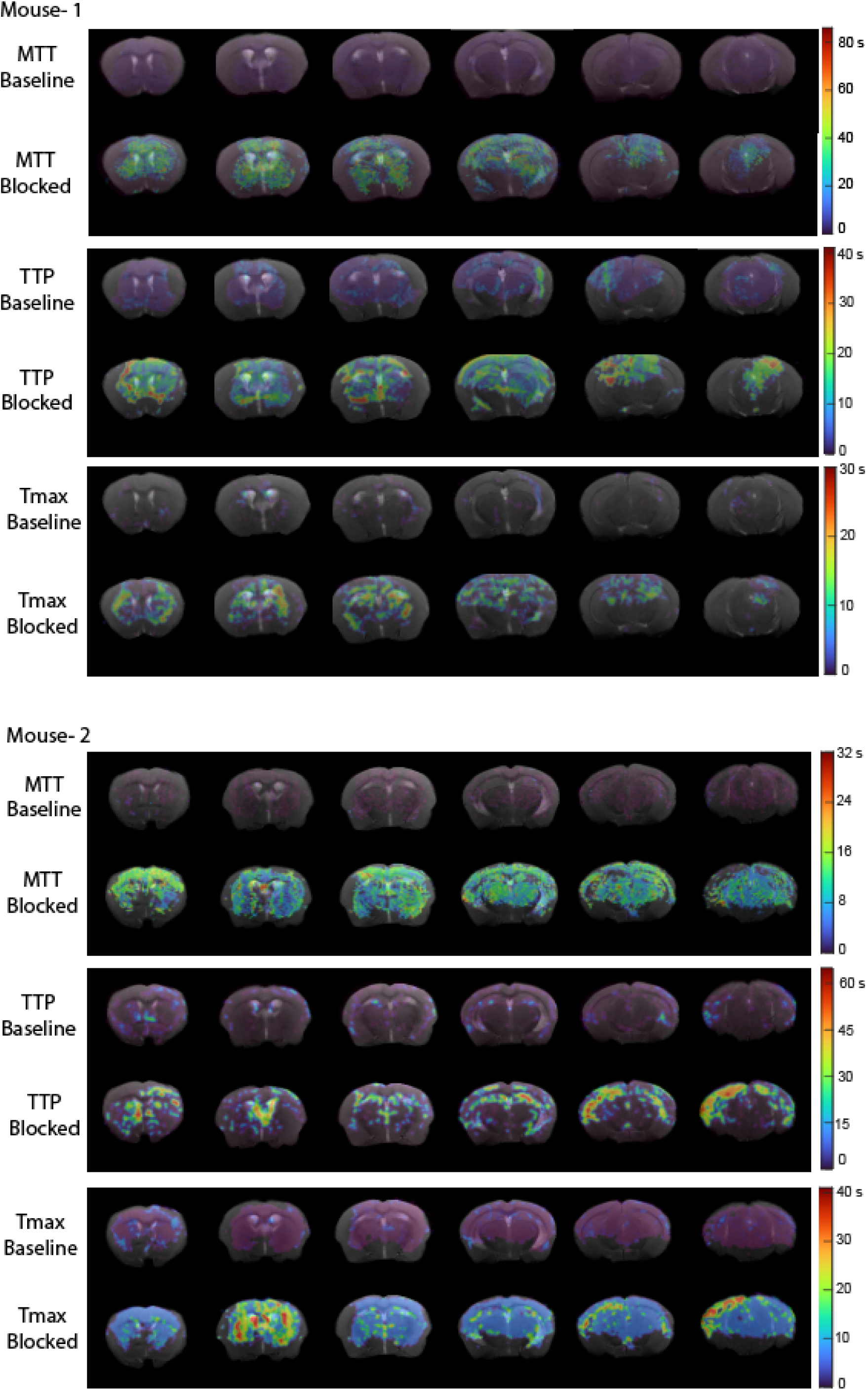
Perfusion maps during micro-balloon–induced CCA occlusion. Representative coronal parametric maps from two mice showing mean transit time (MTT), time-to-peak (TTP), and time-to-maximum (Tmax) before, during, and after micro-balloon occlusion of the CCA. Maps are overlaid on corresponding T2-weighted anatomical images. Colour bars to the right indicate the quantitative range for each parameter. Warmer colours denote longer transit or delay times; cooler colours denote shorter times.

**Extended Data Fig. 18.**
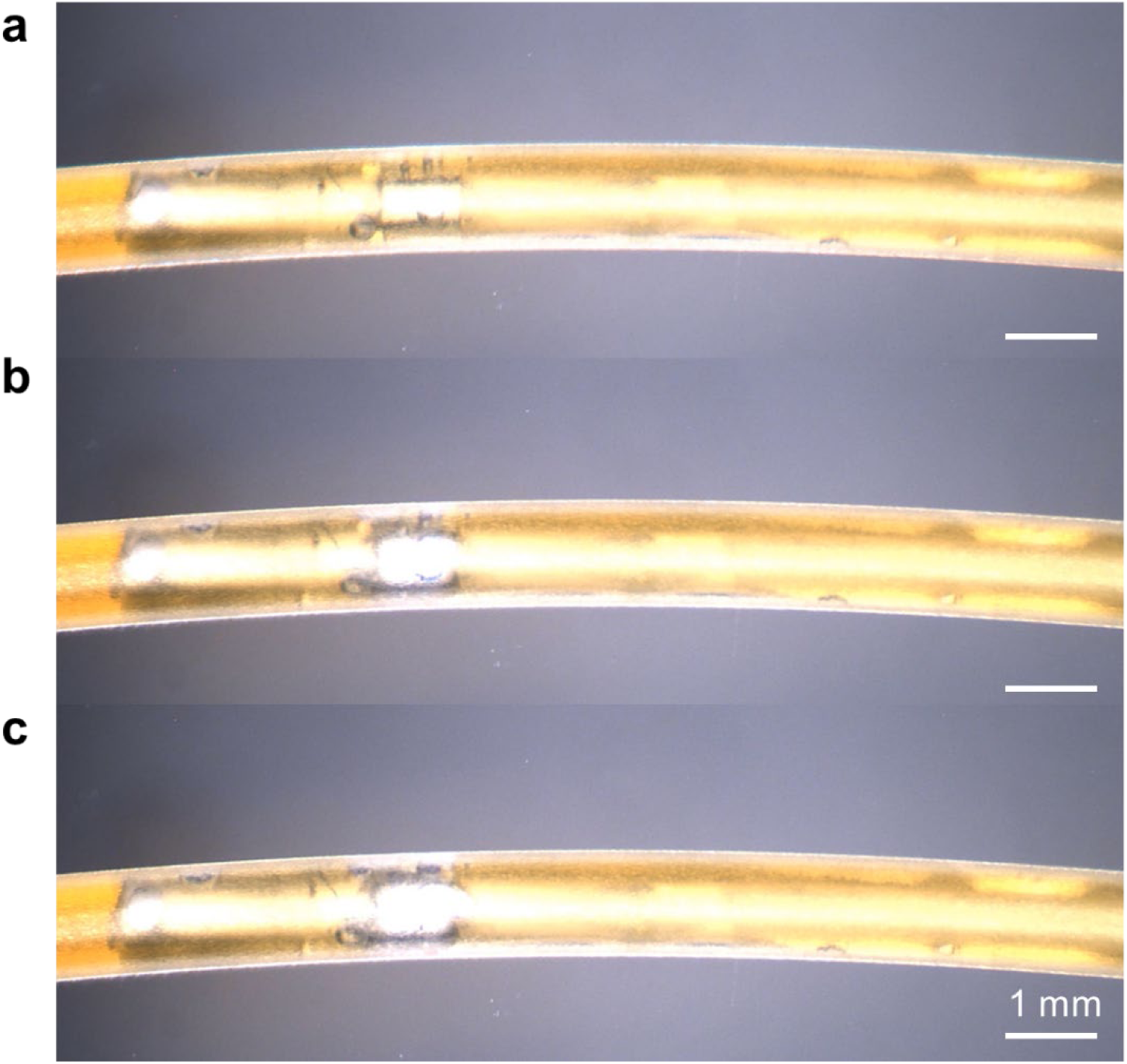
OM images of the micro-balloons blocking water flow inside the plastic tubing: the initial (a), partially blocked (b), and complete blocked (c) states.

**Extended Data Fig. 19.**
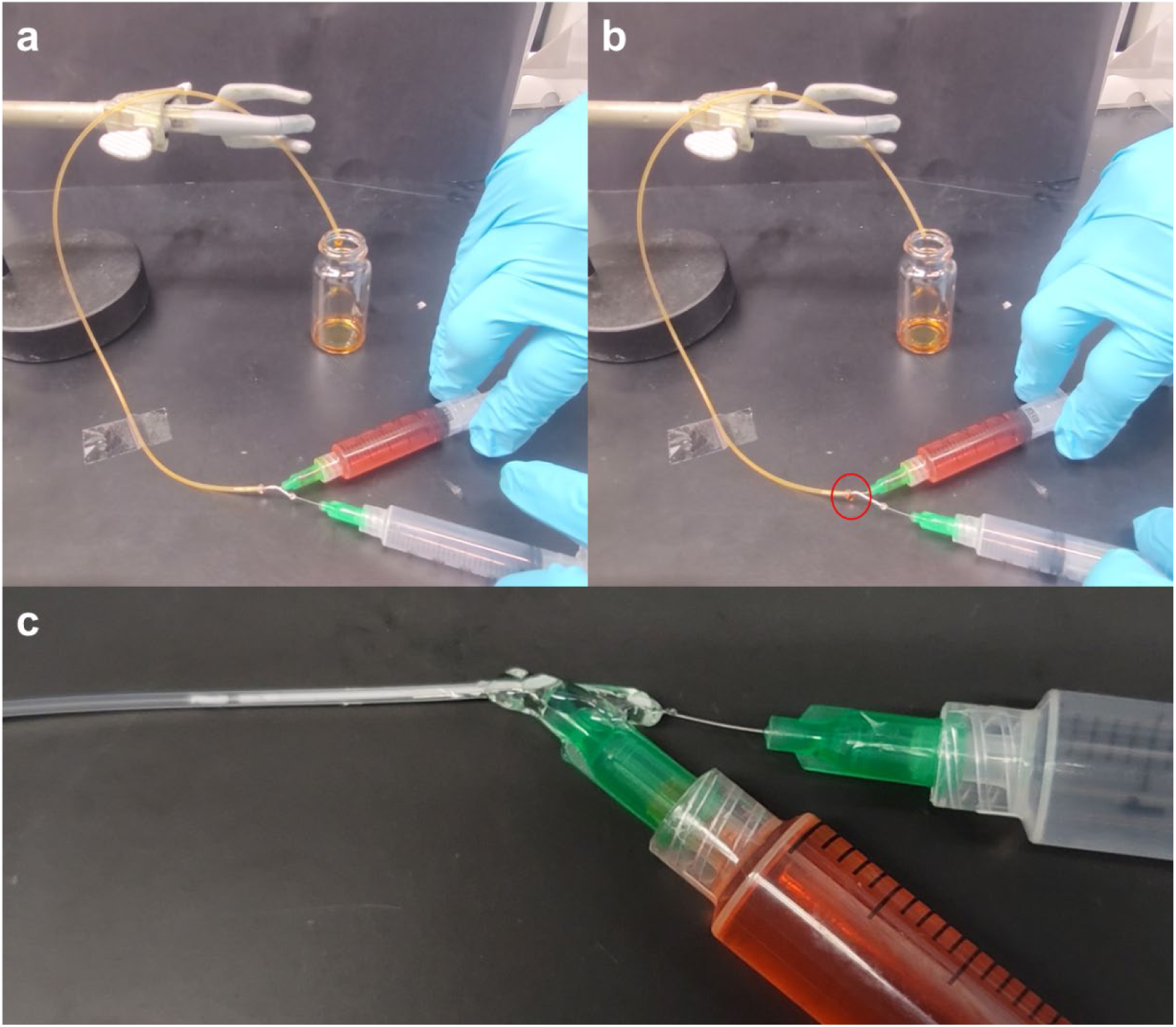
Experimental setup for the intravascular occlusion tests. **a–c,** Before (a) and after (b) pneumatic balloon inflation during water flow, and the magnified view showing the positions of the pneumatic and water-injecting nozzles inside the tubing (c).

**Extended Data Fig. 20.**
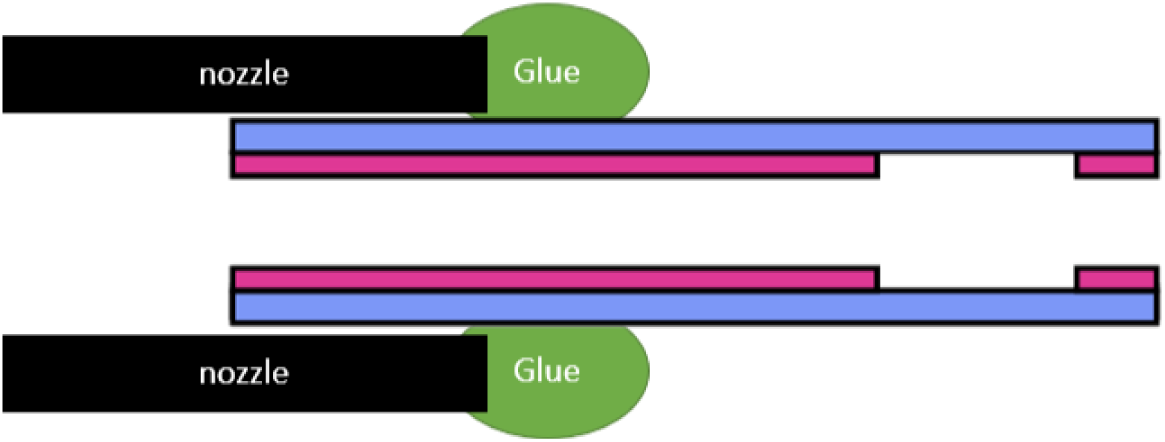
Schematic illustration of how to connect the micro-balloon with an outer diameter of 550 μm to the syringe nozzle.

**Extended Data Table 1.**

|  | **Takeru** | **Abbott** | **Rosalia *et al.***<br>**(*Sci. Robot.*)** | **This work** |
| --- | --- | --- | --- | --- |
| **Balloon initial diameter** | ~ 560 $\mu\text{m}$ | ~ 810 $\mu\text{m}$ | 1.6 mm | ~ 550 $\mu\text{m}$ |
| **Shaft diameter** | ~ 430 $\mu\text{m}$ | ~ 530 $\mu\text{m}$ | 2.4 mm | ~ 550 $\mu\text{m}$ |
| **Balloon length** | ~ 6 mm | ~ 6 mm | 2 mm | ~ 1 mm |
| **Intermediate inflation** | No | No | Yes | Yes |
| **Pressure required** | 6-8 atm | 8 atm | 1~2.33 atm | 1~3 atm |
| **Metal inclusion** | Yes | Yes | No | No |

